# Phosphoinositide regulation by the CCC complex promotes phagosome maturation and host defense

**DOI:** 10.64898/2025.12.15.694371

**Authors:** Hannes Buck, Qi Liu, Amika Singla, Daniel A. Kramer, Rodrigo Pereyria, Wan-Ru Lee, Wesley Burford, Jianyi Yin, Ariel Davis, Shuai Tan, Baoyu Chen, Luis Sifuentes-Dominguez, Neal Alto, Jen Liou, Ezra Burstein

**Affiliations:** Division of Digestive and Liver Diseases, Department of Internal Medicine, University of Texas Southwestern Medical Center, Dallas, TX, USA; Department of Physiology, University of Texas Southwestern Medical Center, Dallas, TX, USA; Department of Microbiology, University of Texas Southwestern Medical Center, Dallas, TX, USA; Department of Biophysics, University of Texas Southwestern Medical Center, Dallas, TX, USA; Department of Pediatrics, University of Texas Southwestern Medical Center, Dallas, TX, USA; Department of Molecular Biology, University of Texas Southwestern Medical Center, Dallas, TX, USA

## Abstract

The evolutionarily conserved COMMD/CCDC22/CCDC93 (CCC) complex regulates endolysosomal function and is required for effective antibacterial immunity. However, the mechanisms linking the CCC complex to macrophage function remain poorly understood. Using bone marrow-derived macrophages, we found that CCC deficiency impaired bacterial clearance and disrupted phagosome maturation, as evidenced by defective phagosomal membrane remodeling, reduced phagosome-endoplasmic reticulum contacts, and impaired phagosome-lysosome fusion. These defects were independent of the CCC complex interaction partner, Retriever. CCC-deficient macrophages exhibited excessive phosphatidylinositol 3-phosphate (PI(3)P) accumulation on phagosomal membranes due to markedly reduced recruitment of the PI(3)P phosphatase MTMR2. Restoration of normal PI(3)P levels rescued bactericidal activity, establishing aberrant PI(3)P regulation as the principal defect affecting CCC-deficient macrophages. These findings identify a previously unrecognized Retriever-independent function of the CCC complex in regulating PI(3)P dynamics during phagosome maturation, a process essential for efficient bacterial clearance and innate immune defense.

## Introduction

Phagocytosis is a fundamental process in innate immune defense whereby cells “ingest” extracellular structures, such as microorganisms, cellular debris, or inorganic materials that may pose a threat to the organism (Li and Underhill, 2025; Stuart and Ezekowitz, 2005). Following engulfment into a large endocytic vesicle known as the phagosome, a variety of degradative enzymes and other effectors are recruited to progressively digest the internalized material. While many mammalian cells are capable of phagocytosis, cells of the myeloid lineage, particularly macrophages and dendritic cells, are highly phagocytic and provide critical host defense through this mechanism (Fu and Harrison, 2021). In addition, as phagosomal contents are degraded, the resulting products can trigger signaling cascades initiating downstream responses, such as cytokine expression and release. The degradation products undergo further processing, resulting in peptides that are loaded onto major histocompatibility complex (MHC) molecules for presentation to T lymphocytes, thereby linking innate and adaptive immunity (Mantegazza et al., 2013).

To accomplish these diverse outcomes, the phagosome undergoes rapid and extensive ultrastructural changes driven by dynamic transitions in its protein and membrane lipid composition (Levin et al., 2015). Immediately following phagosome formation, the phosphoinositide composition of the phagosomal membrane closely resembles that of the plasma membrane (PM), predominantly containing phosphatidylinositol-4-phosphate (PI(4)P), as well as PI(4,5)P_2_ and PI(3,4,5)P_3_ (Levin et al., 2017). Within minutes, this lipid composition is remodeled, and the phagosomal membrane becomes enriched in PI(3)P, a key to the recruitment of multiple effector proteins which are typically associated with early endosomes (Araki et al., 1996; Marshall et al., 2001). During later stages of phagosome maturation, PI(3)P is progressively replaced by PI(3,5)P_2_ and PI(4)P, lipid species that facilitate lysosomal docking and fusion with the phagosomal membrane (Jeschke and Haas, 2018; Jeschke et al., 2015; Kim et al., 2014; Levin et al., 2017; Vines et al., 2023). This stage is pivotal to phagosome maturation, as it is characterized by the delivery of degradative enzymes and antimicrobial factors into the phagosomal lumen, enabling efficient destruction of invading microorganisms. While the recruitment of PI(3)P effectors to phagosomal membranes has been extensively characterized, the mechanisms governing PI(3)P turnover and its precise role in phagosomal maturation remain unclear.

Early phagosomes exhibit significant overlap in both protein and lipid composition with early endosomes (Li and Underhill, 2025). Like early phagosomes, early endosomes are predominately enriched in PI(3)P and serve as key sites for active sorting and recycling of PM proteins. Endosomal recycling is initiated through the recognition of cargo proteins destined for recycling by the evolutionarily conserved sorting complexes Retriever and Retromer (Chen et al., 2019). Following recognition, cargo proteins are segregated into specialized endosomal microdomains containing the COMMD / CCDC22 / CCDC93 (CCC) and WASH complexes (Phillips-Krawczak et al., 2015), two co-evolved assemblies that are indispensable for endosomal recycling. The CCC complex is composed of a heterodimer of CCDC22 and CCDC93, which forms an extended scaffold that, at one end, binds to all ten members of the COMMD protein family assembled into a decameric ring structure. At the opposite end of the CCDC22/CCDC93 scaffold, the CCC complex binds to the Retriever complex, together forming a larger assembly referred to as the Commander complex.

Although the CCC and Retriever complexes cooperate in endosomal protein recycling, whether the CCC complex has functions independent of the Retriever complex remains an outstanding question in the field. This distinction may be particularly relevant in innate immunity. Components of the CCC complex have been reported to localize to phagosomes (Dill et al., 2015; Miyata et al., 2018), and macrophage-specific deletion of *Commd* genes increases susceptibility to sepsis (Ben Shlomo et al., 2019; Cohen et al., 2021; Miyata et al., 2018; Mouhadeb et al., 2018) and elevates bacterial burden within the intestinal lamina propria of mice (Miyata et al., 2018). However, it remains unclear whether these phenotypes are mediated by the CCC complex itself or require the broader Commander complex. In particular, whether other Commander subcomplexes, such as Retriever, contribute to antimicrobial defense has not been investigated. Furthermore, the molecular mechanisms linking the CCC complex to macrophage antibacterial activity remain poorly defined. Here, we address these questions and show that the CCC complex, but not Retriever, is required for efficient phagosome maturation through regulation of PI(3)P turnover.

## Results

### CCC, but not Retriever, is required for E. coli elimination in macrophages

While prior studies have implicated the CCC complex in phagocyte innate immune defense, whether the closely associated Retriever complex exerts similar functions in macrophages has not been previously studied. To examine this question, we generated mice carrying conditional alleles for 2 subunits of the Retriever complex, *Vps35l* and *Vps26c* (**Extended Data Figure 1**). Bone marrow derived macrophages (BMDMs) from *Vps35l* and *Vps26c* knockout (KO) mice demonstrated the expected loss of the targeted protein, but CCC subunits remained affected; in contrast, deletion of *Commd1* or *Commd9* in BMDMs destabilized several components of CCC and Retriever (**Figure 1a**), in agreement with other observations in fibroblasts and epithelial cells (Fedoseienko et al., 2018; Phillips-Krawczak et al., 2015; Singla et al., 2021; Singla et al., 2019). Next, we assessed the impact of such gene deletions on the ability of macrophages to clear commensal non-pathogenic bacteria. Following exposure to live *E. coli*, the intracellular bacterial burden was determined over the subsequent 180 minutes by lysing BMDMs and plating the lysates on antibiotic-containing agar plates to quantify viable bacterial colony-forming units (CFU) the following day. This analysis revealed that wild-type BMDMs achieved over 90% bacterial clearance within the 3 hours of bacterial exposure, whereas *Commd1* KO BMDMs failed to effectively eliminate intracellular *E. coli* (**Figure 1b, Extended Data Figure 2a**). In contrast, *Vps35l* and *Vps26c* KO BMDMs displayed normal bactericidal activity, revealing that Commd1, but not the Retriever subunits Vps35l or Vps26c, contributes to the ability of macrophages to eliminate *E. coli*.

**Figure 1:**
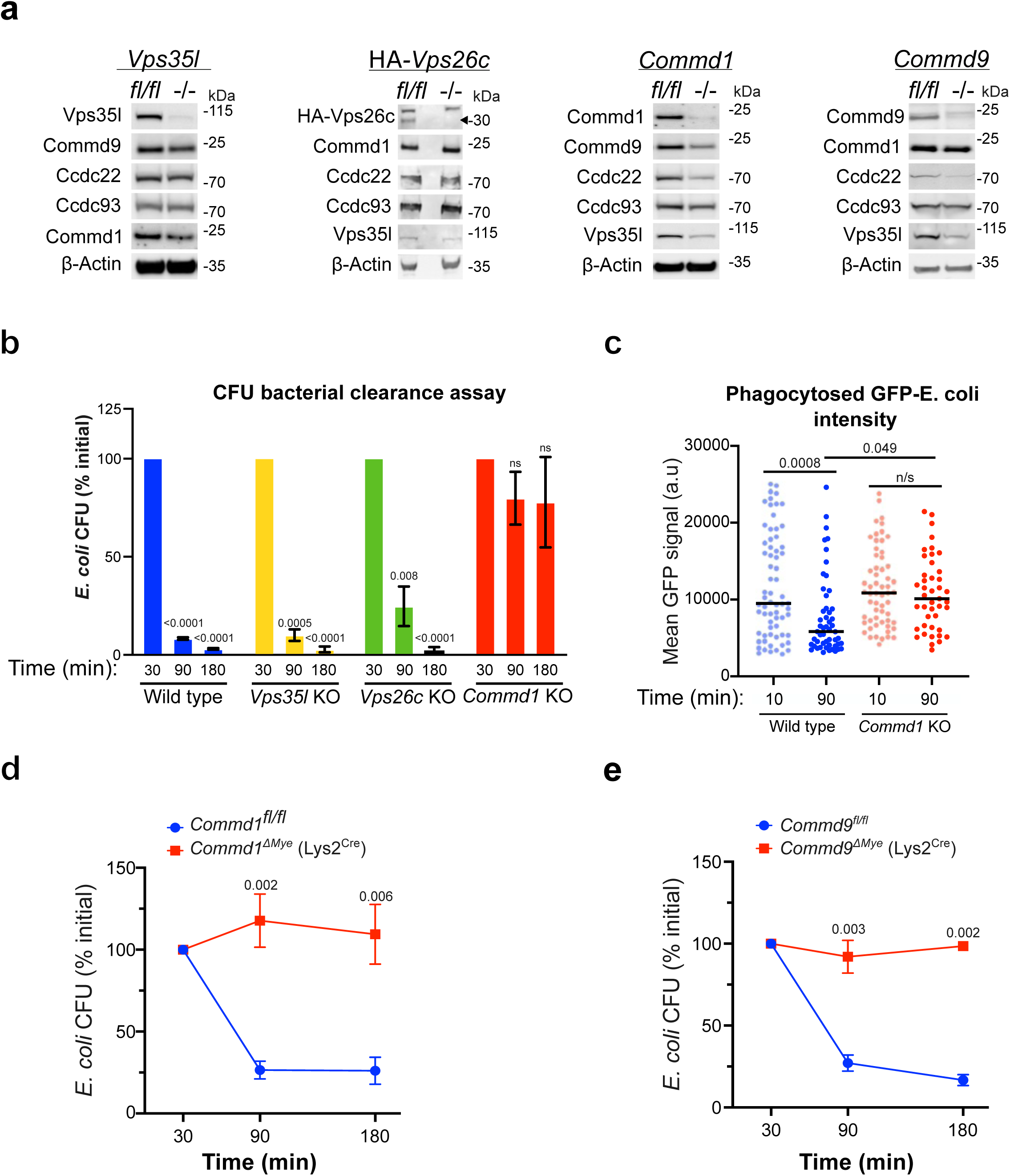
The CCC complex, but not Retriever, is required for intracellular *E. coli* clearance by macrophages. (**a**) Immunoblot analysis of CCC and Retriever complex components in bone marrow–derived macrophages (BMDMs) from wild-type and knockout (KO) mice lacking *Vps35l, Vps26c, Commd1* and *Commd9*. A representative immunoblot from ≥2 independent mice per genotype is shown. The *Vps26c* conditional allele includes the knock-in of an HA tag to allow for antibody detection (see Extended Data Fig 1 for more details). (**b**) Quantification of intracellular bacterial clearance by colony-forming unit (CFU) assay in wild-type, *Commd1* KO, *Vps35l* KO, and *Vps26c* KO BMDMs. Data are representative of at least two independent experiments performed using BMDMs derived from two mice per genotype. (**c**) Quantification of GFP fluorescence intensity from three-dimensionally reconstructed phagocytosed *E. coli* in wild-type and *Commd1* KO BMDMs at the indicated time points. Each dot represents a single bacterium; horizontal bars indicate the mean. Data are representative of two independent imaging experiments, using BMDMs derived from two mice per genotype. (**d**, **e**) Time-course analysis of viable intracellular *E. coli*, measured by CFU assays, in wild-type and the indicated KO BMDMs. Data are representative of five (**d**) or two (**e**) independent experiments performed using BMDMs derived from three mice per genotype. Statistical analysis was performed using one-way ANOVA for multiple-group comparisons in (**b**, **c**) and two-tailed Student’s t-tests for pairwise comparisons between genotypes at each time point in (**d**, **e**).

Next, we confirmed the defect seen in *Commd1* KO macrophages using complementary approaches. BMDMs were plated in a 96-well plate and incubated with GFP-expressing *E. coli*, allowing for phagocytosis over a 30-minute period. After removal of *E. coli* from the media, GFP signal intensity in the well was measured and plotted over time. Unlike wild-type BMDMs that displayed progressive decrease in GFP fluorescence intensity, *Commd1* KO BMDMs failed to reduce GFP signal over the course of a 3-hour period (**Extended Data Figure 2b**). To confirm that this reduction in GFP signal intensity represented loss of bacterial fluorescence inside of phagosomes, we performed confocal imaging after pulsing BMDMs with GFP-expressing *E. coli.* Using three-dimensional reconstruction of intracellular bacteria, we determined GFP signal intensity at early (10 minutes) and late stages (90 minutes) following phagocytosis. This analysis demonstrated that the GFP signal of phagocytosed *E. coli* decreased significantly in wild-type BMDMs but not in *Commd1* KO BMDMs over the course of a 90-minute experiment (**Figure 1c**). We further confirmed the observed results using a different approach to target the Commd1 conditional allele. Whereas the experiments in Figure 2b relied on the *Ubc^CreERT2^* allele (which allows for post-natal tamoxifen-inducible deletion in the entire animal), we generated BMDMs from mice carrying the myeloid-specific and constitutive *Lys2^Cre^* allele. Again, we observed a marked decrease in bactericidal activity in *Commd1* KO BMDMs, confirming that the phenotype is independent of the gene targeting strategy (**Figure 1d**). Lastly, we examined whether this defect was specific to *Commd1* deficiency or, more broadly, the result of CCC complex depletion. We performed the same CFU analysis using *Commd9* KO BMDMs and found that *Commd9* deficiency was associated with a comparable defect in bacterial clearance within the 3-hour window of the experiment (**Figure 1e**). Thus, deficiency in CCC, but not Retriever, results in significant impairment in *E. coli* elimination in BMDMs.

**Figure 2:**
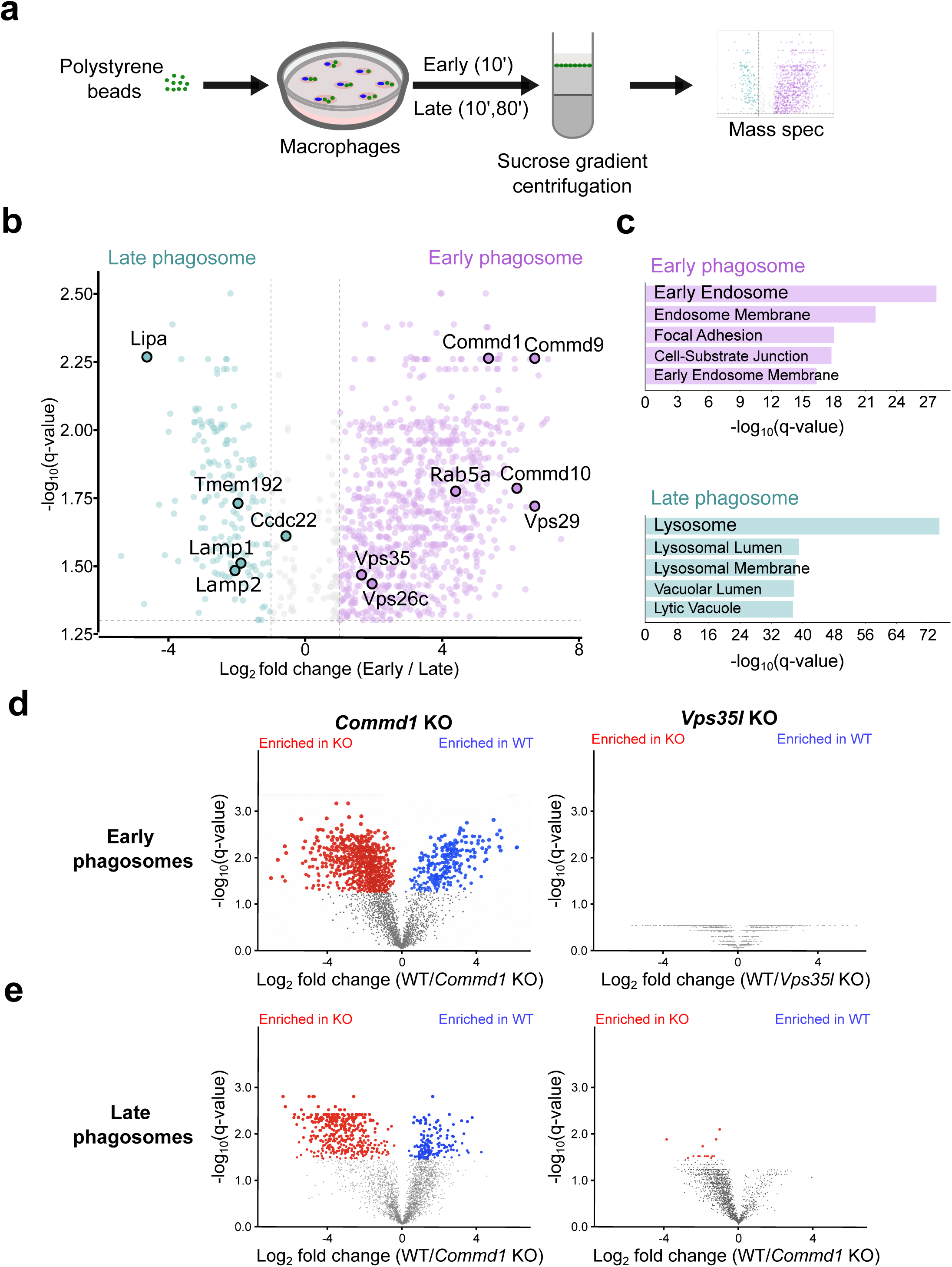
The CCC complex regulates phagosomal protein composition. **(a)** Schematic illustrating phagosome isolation workflow. BMDMs were allowed to internalize polystyrene beads, after which phagosomes were isolated by sucrose-gradient centrifugation and analyzed by mass spectrometry-based proteomics. (**b**) Volcano plot comparing protein abundance between early and late phagosomes. Log_2_ fold change (x axis) is plotted against the FDR-corrected *P* value (y-axis). Data were obtained from three biological replicates using BMDMs derived from three individual wild-type mice. Components of the CCC complex and established markers of early and late phagosomes are highlighted. (**c**) Enrichr analysis of proteins differentially abundant between early and late phagosomes. The top enriched cellular component terms associated with proteins enriched in early phagosomes (top) and late phagosomes (bottom) are shown. (**d**) Volcano plots from independent proteomic analyses comparing early phagosomes isolated from wild-type and *Commd1* knockout BMDMs (left) and from wild-type and *Vps35l* knockout BMDMs (right). (**e**) Volcano plots from independent proteomic analyses comparing late phagosomes isolated from wild-type and *Commd1* knockout BMDMs (left) and from wild-type and *Vps35l* knockout BMDMs (right). Data were obtained from three biological replicates per condition using BMDMs derived from three individual mice per genotype.

### CCC regulates protein composition during phagosome maturation

Phagosomes undergo significant changes in protein and lipid composition over time, a process termed “phagosome maturation” (Lee et al., 2020). Given previous reports that CCC complex subunits are recruited to phagosomes (Dill et al., 2015; Miyata et al., 2018), we hypothesized that the CCC complex may regulate protein composition during phagosome maturation. To examine the potential roles of the CCC and Retriever complexes in phagosome maturation at the proteome level, we isolated phagosomes from BMDMs (**Figure 2a**). Using polystyrene beads as a model phagocytic cargo, analogous to inorganic particulates engulfed by macrophages, we leveraged the low density of the beads to allow phagosome purification by sucrose gradient centrifugation. Isolated phagosomes were subjected to quantitative proteomic analysis to define the protein composition of early and late phagosomes. Changes in protein abundance and their statistical significance were visualized in volcano plots (**Figure 2b**). Consistent with previous reports (Fratti et al., 2001; Vieira et al., 2003), early phagosomes from wild-type BMDMs (upper right quadrant of the volcano plot) were enriched for Rab5a, Retriever complex subunits (Vps26c and Vps29), CCC complex subunits (Commd1, Commd9, and Commd10), and Retromer complex subunits (Vps35 and Vps29). In contrast, late phagosomes from wild-type BMDMs (upper left quadrant of the volcano plot) were enriched in lysosomal proteins (Lipa, Tmem192, Lamp1, and Lamp2), consistent with phagosome fusion with lysosomes at later stages of maturation. Gene set enrichment analysis of proteins preferentially enriched in early phagosomes revealed a profile consistent with that of early endosomes, a closely related organelle (**Figure 2c, top**). In contrast, proteins enriched in late phagosomes showed a strong association with lysosomes, likely reflecting the occurrence of phagosome-lysosome fusion prior to sample collection (**Figure 2c, bottom**).

Immunofluorescence staining for Commd1, Ccdc22, and Ccdc93 in BMDMs exposed to *E. coli* for 10 minutes confirmed the recruitment of these CCC complex subunits to phagosomes (**Extended Data Figure 3**). In contrast to their extensive co-localization on endosomes (Phillips-Krawczak et al., 2015; Singla et al., 2019), these proteins exhibited only partial colocalization on phagosomes, with certain areas of the phagosome membrane decorated by Commd proteins compared to Ccdc22 (top) or Ccdc93 (bottom). Interestingly, the proteomic analysis also revealed distinct recruitment dynamics for these proteins, with Ccdc22 displaying modest enrichment in late phagosomes, whereas Commd proteins were preferentially enriched in early phagosomes (**Figure 2b**). Together, these findings demonstrate that the CCC complex, along with its associated Retriever and Retromer complexes, is recruited to phagosomes during phagosomal maturation.

Next, we evaluated the impact of *Commd1* or *Vps35l* deficiency on phagosomal protein composition of using mass spectrometry-based quantitative proteomics, as described above (**Supplementary Tables 1 and 2**). The proteomic profiles of both early and late phagosomes isolated from *Commd1*-deficient macrophages differed substantially from those of wild-type controls, as shown by volcano plot analysis (**Figures 2d, 2e, left panels**). In contrast, *Vps35l* deficiency had minimal effects in similar experimental settings (**Figures 2d, 2e, right panels, Extended Data Figure 4a**). To validate this observation, we examined whether phagosomal maturation proceeded normally in wild-type macrophages used in both experimental settings. We compared early and late phagosome proteomes from the two wild-type control genotypes (*Commd1^fl/fl^*and *Vps35l^fl/fl^* macrophages) and found that phagosome maturation was evident and comparable in each dataset (**Extended Data Figure 4b, 4c**). Altogether, these findings reveal a profound defect in proteome-level phagosome maturation in the absence of the CCC complex, whereas Retriever deficiency has little detectable impact on this process.

### CCC regulates lysosomal and ER protein recruitment to phagosomes

To determine how loss of *Commd1* alters phagosome maturation, we next examined the functional identity of proteins differentially represented in phagosomes from wild-type and *Commd1*-deficient BMDMs. Pairwise comparison of early phagosomes revealed that proteins normally associated with lysosomes, including Ctsb, Ctsd, Atp6v1a, Tmem192, Lamp1 and Lamp2, were readily detected in early phagosomes from wild-type BMDMs but were significantly underrepresented in phagosomes from *Commd1* KO macrophages (**Figures 3a, 3b**). In contrast, proteins associated with extracellular vesicles and exosomes (e.g., Appl2, A2m, Hps90ab1, and Anxa7) and early endosomes (e.g., Eea1, Vps35, Rab5c, and Snx3) were enriched in early phagosomes from *Commd1*-deficient BMDMs relative to wild-type controls (**Figure 3a, 3c**). Striking differences were also evident in late phagosomes. Compared with *Commd1*-deficient macrophages, wild-type phagosomes had higher abundance of endoplasmic reticulum (ER)-associated proteins (e.g., Vps13c, Sacm1l, and Osbp), including proteins implicated in the formation and function of ER contact sites (Lim et al., 2019; Wang et al., 2025), as well as lysosome-associated proteins (e.g., Vrk2, Fuca1, Npc1, Cln5) (**Figure 3d, 3e**). In contrast, late phagosomes from *Commd1*-deficient macrophages were enriched in proteins associated with extracellular vesicles and exosomes (e.g., Cd9, Cd81, B2m, Anxa5), as well as focal adhesions (e.g., Cdc42, Rac1, Rac2, Rhog) (**Figure 3d, 3f**). Collectively, these findings indicate that *Commd1* deficiency delays phagosome maturation, resulting in prolonged retention of early endosomal and impaired acquisition of lysosomal proteins as well as ER–phagosome contact site-associated proteins.

**Figure 3:**
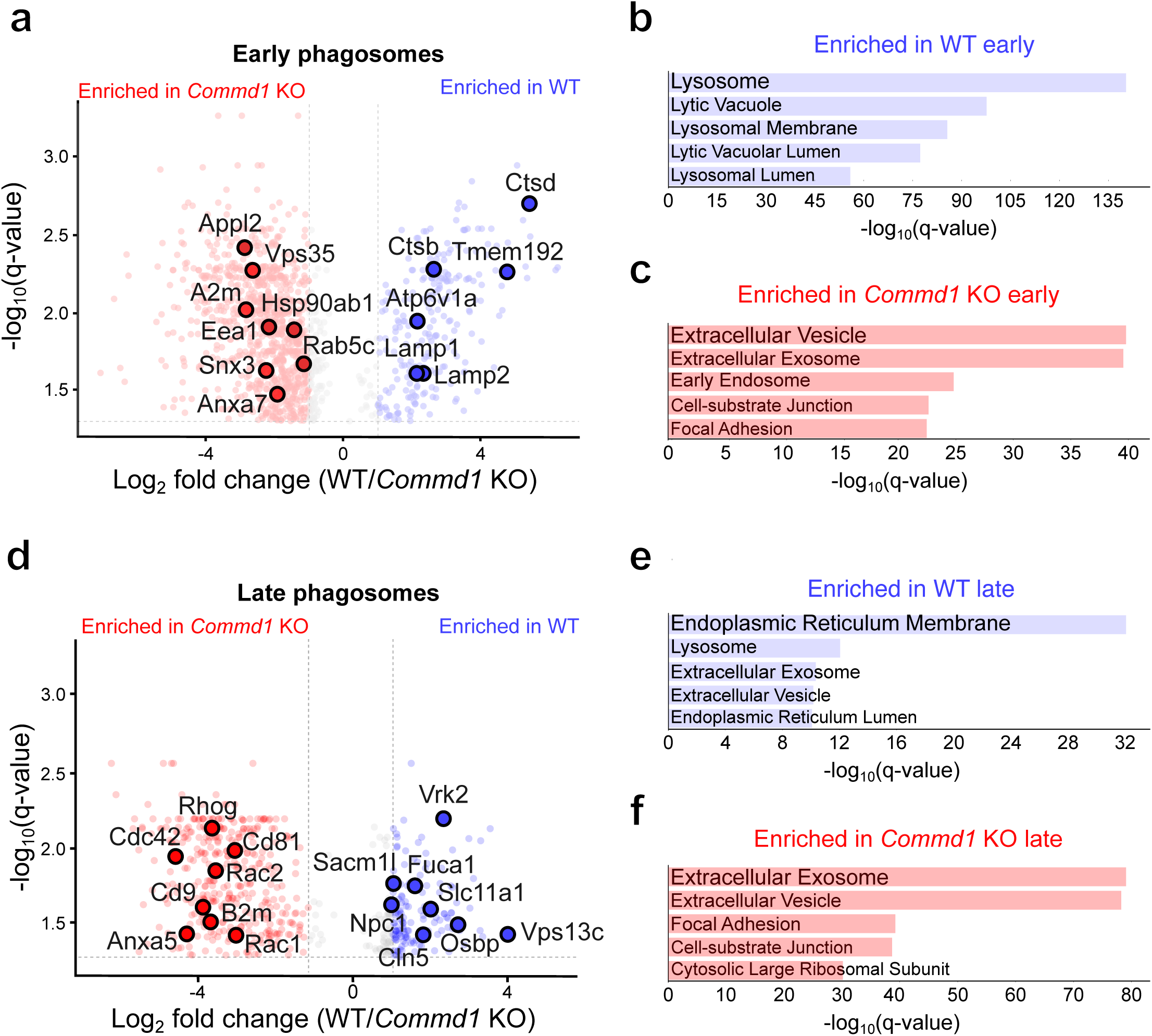
The CCC complex regulates the recruitment of lysosomal and ER-associated proteins to phagosomes. (**a**, **d**) Volcano plots showing proteins identified by mass spectrometry in phagosomes isolated from wild-type and *Commd1* KO BMDMs at early (**a**) and late (**d**) time points. Log_2_ fold change (x-axis) is plotted against -log_10_ of the q-value (FDR-corrected *P* value, y-axis). Highlighted proteins include selected proteins identified through pathway enrichment analyses. (**b**, **c**) Pathway enrichment analyses of proteins differentially abundant in early phagosomes in wild-type (WT) or *Commd1* KO BMDMs, as indicated. (**e**, **f**) Pathway enrichment analyses of proteins differentially abundant in late phagosomes from WT or *Commd1* KO BMDMs, as indicated.

In contrast, early phagosomes from *Vps35l* KO BMDMs minimal alteration compared to wild-type macrophages (**Figure 2d**), whereas late phagosomes displayed only modest changes, primarily involving PM-associated proteins (**Figure 2e, Extended Data Figure 4a**). These changes likely reflect defects in membrane protein recycling resulting from Retriever deficiency (McNally et al., 2017; Singla et al., 2024; Singla et al., 2019). Together, these data indicate that the CCC complex, but not Retriever, is required for the acquisition of key components involved in phagosome maturation. These findings further suggest that phagolysosome formation, a critical step in the degradation of internalized cargo through phagosome fusion with lysosomes, is compromised in the absence of the CCC complex.

### The CCC complex is required for phagosome-lysosome fusion

To validate the proteomic evidence for impaired lysosomal protein acquisition, we examined phagosome-lysosome fusion using complementary biochemical and imaging approaches. First, recruitment of Lamp1, a lysosome-enriched membrane protein, to phagosomes was assessed by flow cytometry. Magnetic beads were used to isolate phagosomes, which were subsequently immunostained for Lamp1 and analyzed by flow cytometry (**Figure 4a, Extended Data Figure 5a, b**). Using this approach, we observed an approximately 8-fold increase in median Lamp1 fluorescence during phagosome maturation in wild-type BMDMs (**Extended Data Figure 5c, 5d**), consistent with progressive phagosome-lysosome fusion. Using this approach, we observed that *Commd1*-deficient macrophages exhibited ∼2-fold reduction in Lamp1 signal on late phagosomes compared with wild-type BMDMs (**Figure 4b, 4c**). We next determined whether lysosomal protein acquisition was similarly impaired during phagocytosis of *E. coli*. Using immunofluorescence microscopy, we assessed the localization of the lysosomal membrane protein Tmem192 to *E. coli*-containing phagosomes (**Figure 4d**). In wild-type BMDMs, Tmem192 staining surrounding internalized bacteria increased over time, reflecting ongoing fusion with lysosomal compartments. In contrast, *Commd1* KO BMDMs failed to show a significant increase in Tmem192 recruitment during the same period (**Figure 4e**).

**Figure 4:**
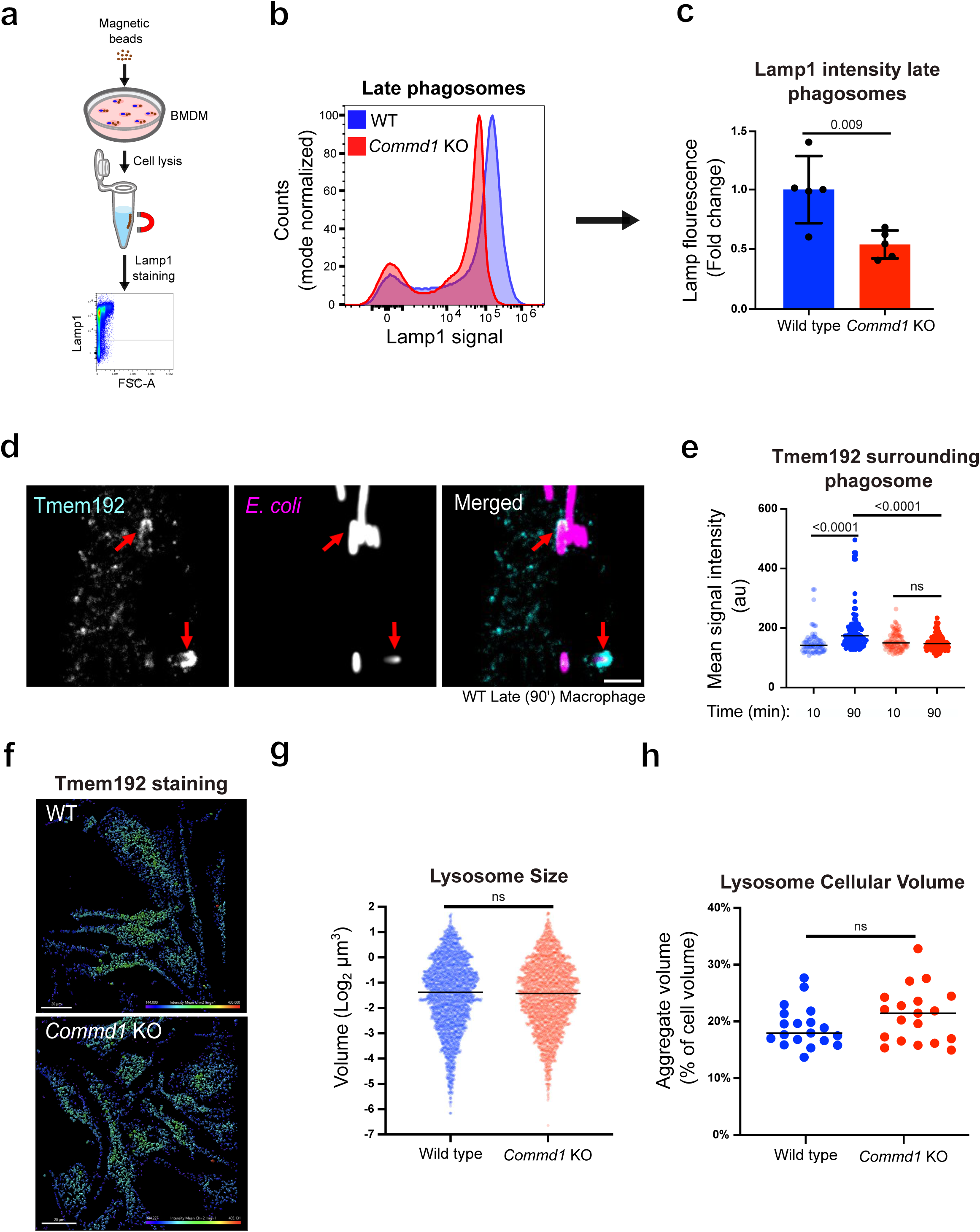
The CCC complex is required for phagosome–lysosome fusion. **(a)** Schematic of the flow cytometry-based workflow used to quantify phagosomal Lamp1 acquisition; isolated phagosomes following magnetic bead uptake were used for immunostaining and flow cytometry. (**b**) Representative histogram of Lamp1 fluorescence on isolated late phagosomes from wild-type and *Commd1* KO BMDMs. (**c**) Quantification of Lamp1 median fluorescence intensity on isolated late phagosomes from wild-type and *Commd1* KO BMDMs. Data are representative of five independent experiments performed using BMDMs derived from three mice per genotype. (**d**) Representative confocal images of GFP-expressing *E. coli*-containing phagosomes in wild-type macrophages at a late stage of maturation. Tmem192 is shown in teal and GFP-expressing *E. coli* in magenta. Arrows indicate Tmem192-positive membranes surrounding bacteria. (**e**) Quantification of Tmem192 fluorescence intensity associated with *E. coli*-containing phagosomes in wild-type and *Commd1* KO macrophages at the indicated time points. Data are representative of two independent experiments using BMDMs derived from two mice per genotype. (**f**) Representative three-dimensional reconstructions of confocal images of macrophages immunostained for Tmem192. Lysosomal structures are rendered according to Tmem192 fluorescence intensity. (**g**) Quantification of Tmem192-positive lysosome volume. Data were obtained from 20 cells and are representative of two independent experiments using BMDMs derived from two mice per genotype. (**h**) Quantification of total lysosomal content per cell, expressed as the percentage of cellular volume occupied by Tmem192-positive structures in three-dimensionally reconstructed cells. Cell boundaries were defined by phalloidin staining. Data were obtained from 20 cells and are representative of two independent experiments using BMDMs derived from two mice per genotype. One-way ANOVA was used for multiple-group comparisons in (**e**). Two-tailed Student’s *t*-tests were used for pairwise comparisons in (**c**), (**g**) and (**h**). Scale bar, 3 μm.

Given the reduced acquisition of lysosomal proteins by phagosomes in *Commd1-*deficient BMDMs, we next investigated whether lysosome abundance or morphology was altered. Unstimulated macrophages were immunostained for Tmem192, and three-dimensional reconstructions were generated from Z-stack images (**Figure 4f**). Lysosome volume (μm^3^) and total lysosomal content per cell, defined as lysosomal volume expressed as a percentage of total cellular volume, were then quantified. Neither measure was significantly affected by *Commd1* deficiency (**Figures 4g, 4h**), indicating that lysosome abundance and morphology remain largely intact in the absence of the CCC complex. Together, these findings demonstrate that the CCC complex is required for efficient delivery of lysosomal proteins to maturing phagosomes. The observed defect occurs without detectable changes in lysosome organelle abundance, suggesting that impaired phagosome maturation results from defective phagosome–lysosome fusion rather than altered lysosome biogenesis. These results prompting us to investigate the ultrastructural basis of the phagosome maturation defect associated with loss of the CCC complex.

### CCC deficiency disrupts ultrastructural maturation of the phagosome

Electron microscopy (EM) enables visualization of key features of phagosome maturation, including ER-phagosome membrane contact sites, progressive bacterial degradation, and increased electron density within the phagosomal lumen following fusion with electron-dense lysosomes. To assess these features, macrophages containing internalized bacteria were fixed and imaged at defined early and late time points of phagosome maturation (**Figure 5a**). This analysis revealed progressive morphological degradation of phagocytosed bacteria in wild-type macrophages, accompanied by a reduction in bacterial electron density, indicative of bacterial destruction within maturing phagosomes (**Figure 5b**). In contrast, these changes were markedly attenuated in *Commd1* KO BMDMs, consistent with the impaired bactericidal activity observed in *Commd1-*deficient macrophages (**Figure 1**). In addition, late phagosomes in wild-type macrophages exhibited greater luminal electron density than early phagosomes, as expected following phagosome-lysosome fusion and the delivery of lysosomal contents (**Figure 5c, 5d**). In contrast, phagosomes in *Commd1*-deficient BMDMs failed to acquire comparable increases in electron density during maturation, consistent with a defect in phagolysosome formation. We also observed that in wild-type macrophages there was a significant increase in the number of lysosomes located within 500 nm of late phagosomes compared to early phagosomes; in stark contrast, this increase was absent in *Commd1* deficient macrophages (**Figure 5g**), indicating impaired lysosome recruitment during phagosome maturation. Another notable finding was the enlarged size of late phagosomes in *Commd1* KO macrophages compared with those in wild-type cells (**Figure 5e)**, suggesting altered phagosomal membrane dynamics, including possible defects in membrane fission. Lastly, our proteomic analyses revealed relative depletion of ER-associated proteins in late phagosomes from *Commd1* KO macrophages. We posited that this represented reduced ER-phagosome contact site formation (Ghavami and Fairn, 2022). Consistent with this prediction, quantitative analyses of EM images revealed a marked reduction in ER-phagosome membrane contact sites in *Commd1* KO macrophages compared with wild-type controls (**Figure 5f, 5h**). These findings corroborate the proteomic data and identify impaired ER-phagosome interactions as an additional consequence of CCC deficiency. Together, these results demonstrate that CCC deficiency disrupts multiple ultrastructural hallmarks of phagosome maturation, including lysosome recruitment, bacterial degradation, lysosomal content acquisition, phagosomal membrane remodeling, and ER–phagosome membrane contact site formation. These findings prompted us to investigate the molecular mechanisms underlying the role of the CCC complex in phagosome maturation.

**Figure 5:**
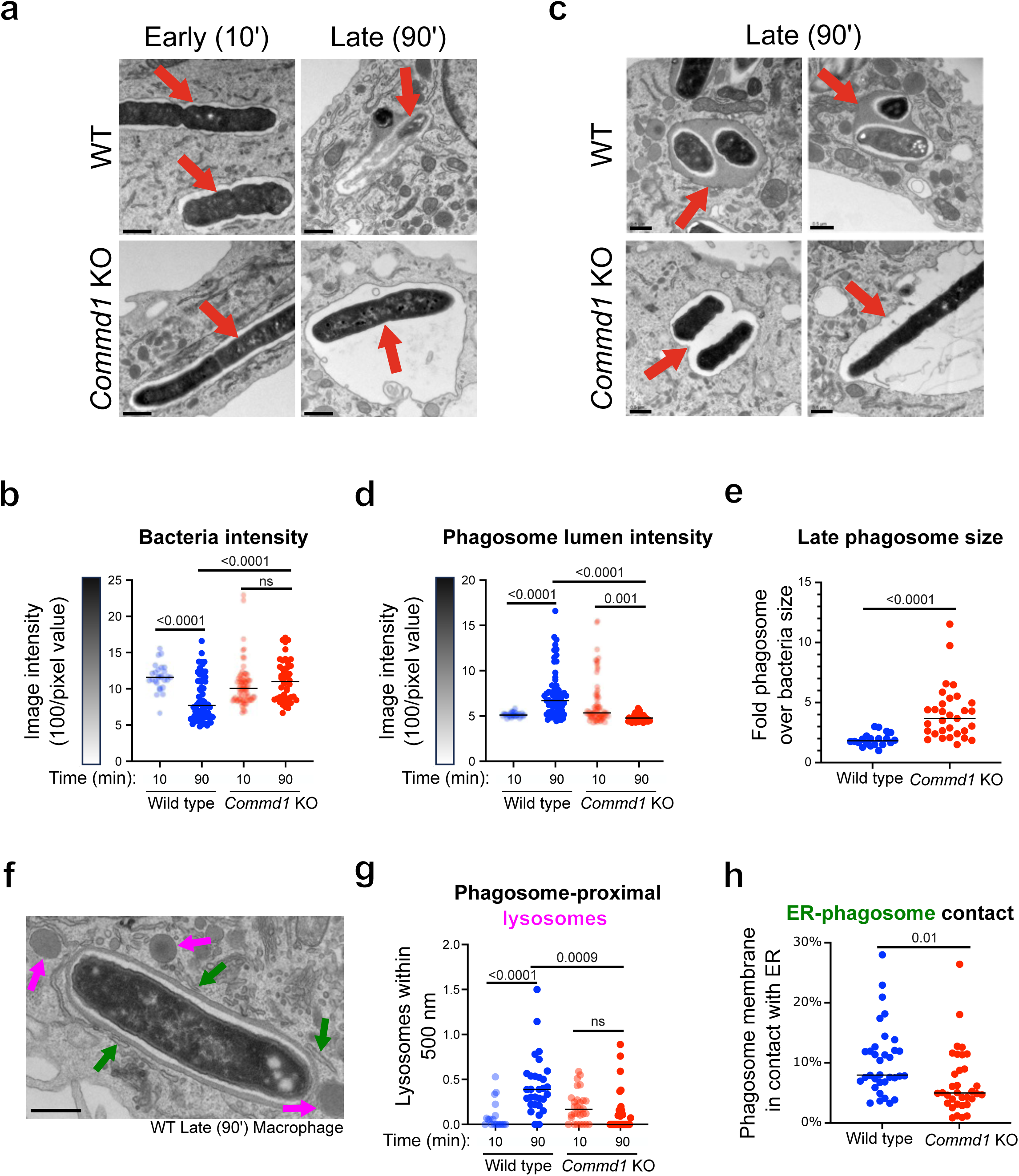
Commd1 deficiency alters ultrastructural features of phagosome maturation. **(a, b)** Representative electron micrographs of early and late phagosomes from wild-type and *Commd1* KO BMDMs. Arrows indicate phagocytosed *E. coli* (**a**) or the phagosomal lumen (**b**). (**c**) Quantification of mean electron density of phagocytosed *E. coli* within phagosomes, measured as mean pixel intensity. (**d**) Quantification of phagosomal lumen electron density measured as mean pixel intensity of the phagosomal space surrounding internalized *E. coli*. (**e**) Quantification of late phagosome size in wild-type and *Commd1* KO BMDMs. (**f**) Representative electron micrograph of a late phagosome from a wild-type macrophage. Green arrows indicate ER–phagosome membrane contact sites, while purple arrows indicate lysosomes. (**g**) Quantification of the number of lysosomes located within 500 nm of phagosomes in the indicated conditions. (**h**) Quantification of ER–phagosome membrane contact sites, expressed as the percentage of the phagosomal membrane closely apposed to the ER. All data presented in this figure is representative of two independent experiments using BMDMs derived from two mice per genotype. One-way ANOVA was used for multiple-group comparisons in (**c**) and (**d**). Two-tailed Student’s *t-*tests were used for pairwise comparisons in (**e**) and (**g**). Scale bars, 0.5 μm.

### The CCC complex controls phagosomal phosphoinositide composition

Selective enrichment of specific PIPs contributes to the establishment of organelle membrane identity and regulates the recruitment of downstream effector proteins (Posor et al., 2022). In addition, PIPs play critical roles in vesicle fusion (Stroupe et al., 2006), as well as the formation and function of ER membrane contact sites with multiple organelles (Raiborg et al., 2016), including phagosomes. Among these lipids, PI(3)P is a hallmark of early endosomes and early phagosomes, where it promotes the recruitment of proteins required for maturation to these vesicles (Birkeland and Stenmark, 2004). Another critical PIP is PI(4)P, which has been implicated in later stages of phagosome maturation and ER-phagosome interactions (Ghavami and Fairn, 2022; Levin et al., 2017). To investigate whether phosphosinositide-dependent pathways were altered in the absence of the CCC complex, we reanalyzed our phagosomal proteomic dataset with a focus on proteins known to bind PI(3)P or PI(4)P. Strikingly, twelve PI(3)P-binding proteins (Gillooly et al., 2001a; Kutateladze, 2006) were significantly enriched in early phagosomes isolated from *Commd1*-deficient BMDMs compared with WT controls (**Figure 6a**). Notably, none of these proteins were significantly altered in *Vps35l*-deficient BMDMs. Conversely, proteins known to bind PI(4)P were significantly depleted from late phagosomes in *Commd1*-deficient macrophages, whereas no comparable changes were observed following *Vps35l* deficiency (**Figure 6b**).

**Figure 6:**
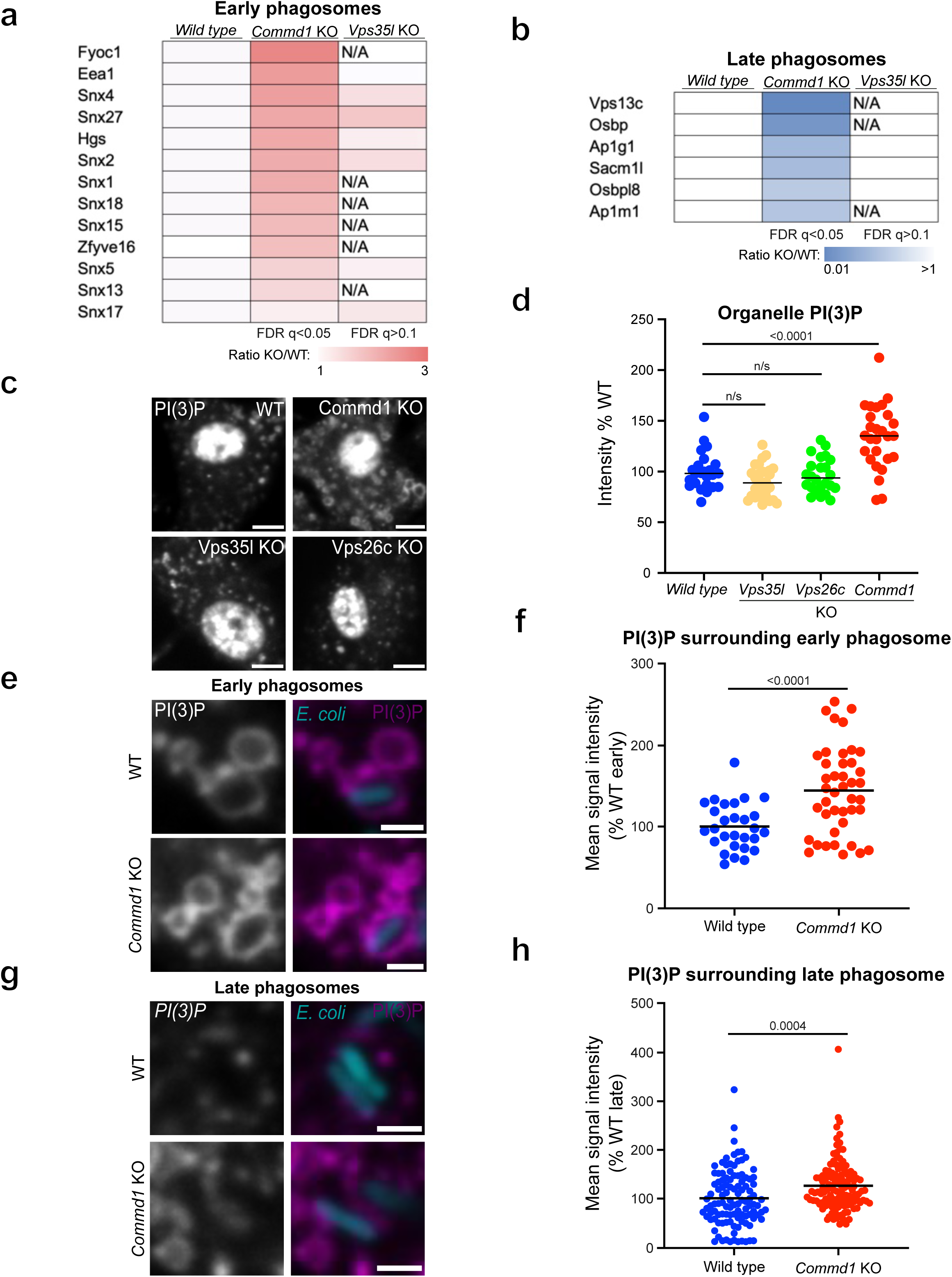
The CCC complex controls phagosomal phosphoinositide composition. **(a)** Heatmap showing the abundance of PI(3)P-binding proteins identified by phagosomal proteomic analyses in early phagosomes isolated from wild-type, *Commd1* KO and *Vps35l* KO BMDMs. (**b**) Heatmap showing the abundance of PI(4)P-binding proteins identified by phagosomal proteomic analyses in late phagosomes isolated from wild-type, *Commd1* KO and *Vps35l* KO BMDMs. (**c**) Representative images of PI(3)P staining in wild-type, *Commd1* KO, *Vps35l* KO and *Vps26c* KO BMDMs. (**d**) Quantification of whole-cell PI(3)P fluorescence intensity, excluding nuclei, in wild-type, *Commd1* KO, *Vps35l* KO and *Vps26c* KO BMDMs. Data are representative of at least two independent experiments from BMDMs isolated from two or more mice per genotype. (**e**) Representative images of PI(3)P-positive early phagosomes (10-min pulse) in wild-type and *Commd1* KO macrophages. (**f**) Quantification of PI(3)P fluorescence intensity surrounding phagocytosed E. coli in wild-type and *Commd1* KO macrophages at the indicated time points. Data are representative of three independent experiments from BMDMs isolated from two mice per genotype. (**g**) Representative images of PI(3)P-positive late phagosomes (10-min pulse, 80-min chase) in wild-type and *Commd1* KO macrophages. (**h**) Quantification of PI(3)P fluorescence intensity surrounding phagocytosed E. coli in wild-type and *Commd1* KO macrophages at the indicated time points. Data are representative of three independent experiments from BMDMs isolated from two mice per genotype. One-way ANOVA was used for multiple-group comparisons in (**d**). Two-tailed Student’s *t*-tests were used for pairwise comparisons in (**f**) and (**h**). Images in (**c**), (**e**) and (**g**) were processed using identical threshold and display settings. Scale bars, 5 μm (**c**) and 2 μm (**e**, **g**).

To directly assess phosphoinositide distribution, we employed a recently described recombinant fluorescent probe-based approach (Maib et al., 2024). Briefly, recombinant proteins containing phosphoinositide-binding domain(s) fused to a SNAP tag were fluorescently labeled *in vitro* and used as staining reagents. We generated probes specific for PI(3)P, consisting of tandem FYVE domains, and PI(4)P, containing the SidC PI(4)P-binding domain. Following saponin permeabilization, these probes were used to stain fixed BMDMs. The PI(3)P probe prominently labeled small intracellular vesicles, likely corresponding to early endosomal compartments (**Figure 6c**). In contrast, the PI(4)P probe strongly labeled the Golgi region but showed minimal signal on endosomes or phagosomes, limiting its utility for subsequent analyses (**Extended Data Figure 7a**).

Quantitative imaging revealed increased cellular PI(3)P abundance in *Commd1* KO macrophages, whereas *Vps35l* KO and *Vps26c* KO macrophages showed no significant changes compared with wild-type cells (**Figure 6d**). Moreover, PI(3)P-positive structures in *Commd1* KO macrophages were significantly enlarged (**Extended Data Figure 7b**). This phenotype is reminiscent of the enlarged late phagosomes observed by EM and is consistent with the endosomal enlargement reported in other settings of CCC complex deficiency (Singla et al., 2019). We next examined PI(3)P localization on *E. coli*-containing phagosomes. Early phagosomes exhibited ring-like PI(3)P staining around the phagosomal membrane, whereas late phagosomes displayed a more punctate pattern (**Figure 6e, 6g**), likely reflecting ongoing PI(3)P turnover during maturation (Gillooly et al., 2001b; Vines et al., 2023). Importantly, *Commd1* deficiency was associated with significantly increased PI(3)P signal surrounding both early and late phagosomes, consistent with the proteomic enrichment of PI(3)P-binding proteins (**Figure 6e-h**). Altogether, these findings demonstrate that loss of the CCC complex results in abnormal accumulation of PI(3)P, both globally and on phagosomal membranes. The specificity of this phenotype to *Commd1* deficiency, together with the associated depletion of PI(4)P-associated proteins from late phagosomes, suggests that the CCC complex plays a critical role in regulating phosphoinositide composition during phagosome maturation.

### CCC deficiency leads to reduced PI(3)P turnover on phagosomes

PI(3)P generation on endosomal and phagosomal membranes is mediated by the Vps34-containing class III phosphoinositide 3-kinase (PI3K) complex (Backer, 2016; Giridharan et al., 2022). In contrast, PI(3)P turnover is driven by lipid phosphatases of the myotubularin (MTMR) family (Dewi Pamungkas Putri et al., 2019; Sheffield et al., 2019; Singla et al., 2019). We therefore reasoned alterations in the enzymes responsible for PI(3)P synthesis or breakdown could account for the abnormal accumulation of PI(3)P observed in *Commd1*-deficient macrophages. When we examined our phagosomal proteomic datasets for evidence of altered kinase of phosphatase recruitment, these proteins were not detected at levels sufficient for reliable quantification, likely reflecting the low abundance of lipid-modifying enzymes required to drive substantial changes in membrane phosphoinositide composition. We therefore employed magnetic bead-based phagosome isolation followed by immunoblotting to directly assess recruitment of PI(3)P regulatory enzymes to phagosomes. The abundance of Vps34 on phagosomes was moderately reduced in *Commd1*-deficient BMDMs (**Figure 7a**), arguing against increased PI(3)P production as the cause of PI(3)P accumulation. In contrast, recruitment of the PI(3)P phosphatase Mtmr2 to phagosomes was drastically reduced in *Commd1*-deficient macrophages (**Figure 7b**). These findings suggest that elevated PI(3)P levels in *Commd1* deficient macrophages result primarily from impaired PI(3)P turnover due to deficient Mtmr2 recruitment.

**Figure 7:**
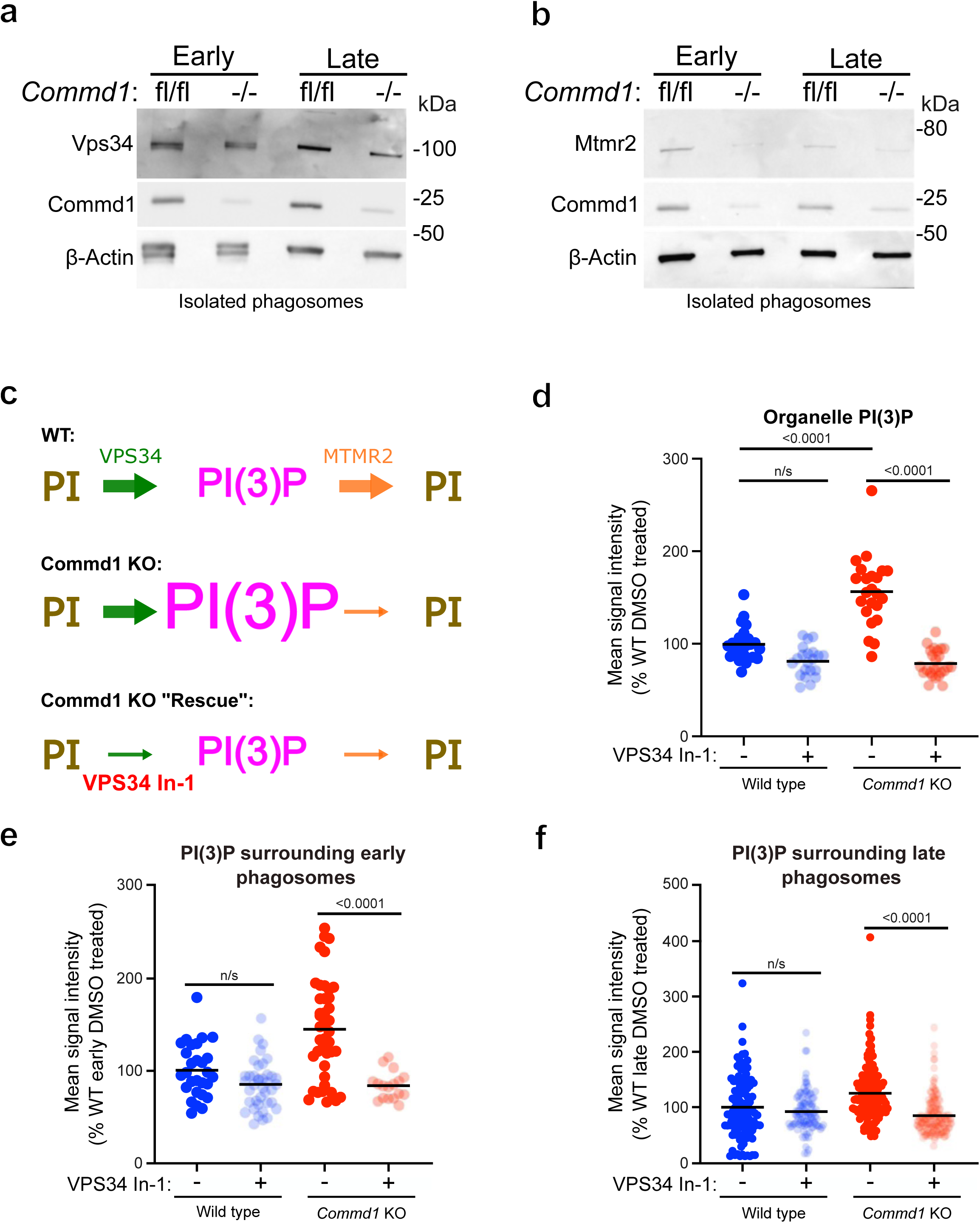
CCC deficiency impairs PI(3)P turnover on phagosomes. (**a**) Immunoblot analysis of Vps34 in isolated phagosomes from wild-type and *Commd1* KO BMDMs. Representative of three independent experiments. (**b**) Immunoblot analysis of Mtmr2 in isolated phagosomes from wild-type and *Commd1* KO BMDMs. Representative of two independent experiments. (**c**) Models illustrating PI(3)P dynamics on phagosomal membranes and the pharmacological strategy used to normalize elevated PI(3)P levels in *Commd1* KO phagosomes through partial inhibition of Vps34. (**d**) Quantification of whole-cell PI(3)P fluorescence intensity, excluding nuclei, in wild-type and *Commd1* KO BMDMs treated with DMSO or VPS34-IN-1. Data are representative of three independent experiments from BMDMs isolated from two mice per genotype. (**e**, **f**) Quantification of PI(3)P fluorescence intensity surrounding phagocytosed *E. coli* in wild-type and *Commd1* KO macrophages treated with DMSO or VPS34-IN-1 at the indicated early (10-min pulse) and late (10-min pulse, 90-min chase) time points. Data are representative of two independent experiments from BMDMs isolated from two mice per genotype. One-way ANOVA was used for multiple-group comparisons in (**d**, **e**, **f**).

### Restoring PI(3)P balance rescues bactericidal activity in CCC-deficient macrophages

The selective accumulation of PI(3)P in CCC-deficient, but not Retriever-deficient, macrophages led us to hypothesize that dysregulated PI(3)P metabolism may be the direct cause for the bactericidal defects observed in *Commd1*-deficient macrophages. To test this hypothesis, we sought to restore PI(3)P homeostasis by reducing PI(3)P synthesis to compensate for the diminished turnover expected from reduced Mtmr2 recruitment (**Figure 7c**). To achieve this, we treated cells with VPS34 In-1, a selective Vps34 inhibitor, at a concentration chosen to partially suppress PI(3)P production while preserving basal Vps34 activity (Bago et al., 2014). Treatment with VPS34 In-1 for 30 minutes normalized the elevated cellular PI(3)P levels observed in *Commd1*-deficient BMDMs (**Figure 7d**). Similarly, the increased PI(3)P signal detected on phagosomal membranes in *Commd1* KO BMDMs was restored to wild-type levels at both early and late stages of phagosome maturation (**Figure 7e, 7f**). Having successfully corrected PI(3)P accumulation, we next examined the impact of this intervention on bactericidal activity. Wild-type macrophages efficiently eliminated *E. coli* irrespective of VPS34-IN-1 treatment, indicating that partial inhibition of Vps34 at the concentrations used did not impair bacterial killing (**Figure 8a**, blue bars). Strikingly, treatment with VPS34-IN-1 completely rescued the bactericidal defect of *Commd1*-deficient macrophages (**Figure 8a**, red bars). Consistent with these findings, 3D reconstructions of macrophages exposed to GFP-expressing *E. coli* demonstrated persistent GFP signal in untreated *Commd1* KO BMDMs at 90 minutes, reflecting defective bacterial clearance. This phenotype was largely reversed following VPS34-IN-1 treatment (**Figure 8b**). Altogether, these results demonstrate that loss of the CCC complex impairs phagosomal PI(3)P turnover through defective recruitment of the PI(3) phosphatase Mtmr2, leading to abnormal PI(3)P accumulation. Importantly, pharmacological restoration of PI(3)P homeostasis fully rescues the bactericidal defect of *Commd1*-deficient macrophages, identifying dysregulated PI(3)P metabolism as a key mechanism underlying defective phagosome maturation and bacterial killing in the absence of the CCC complex.

**Figure 8:**
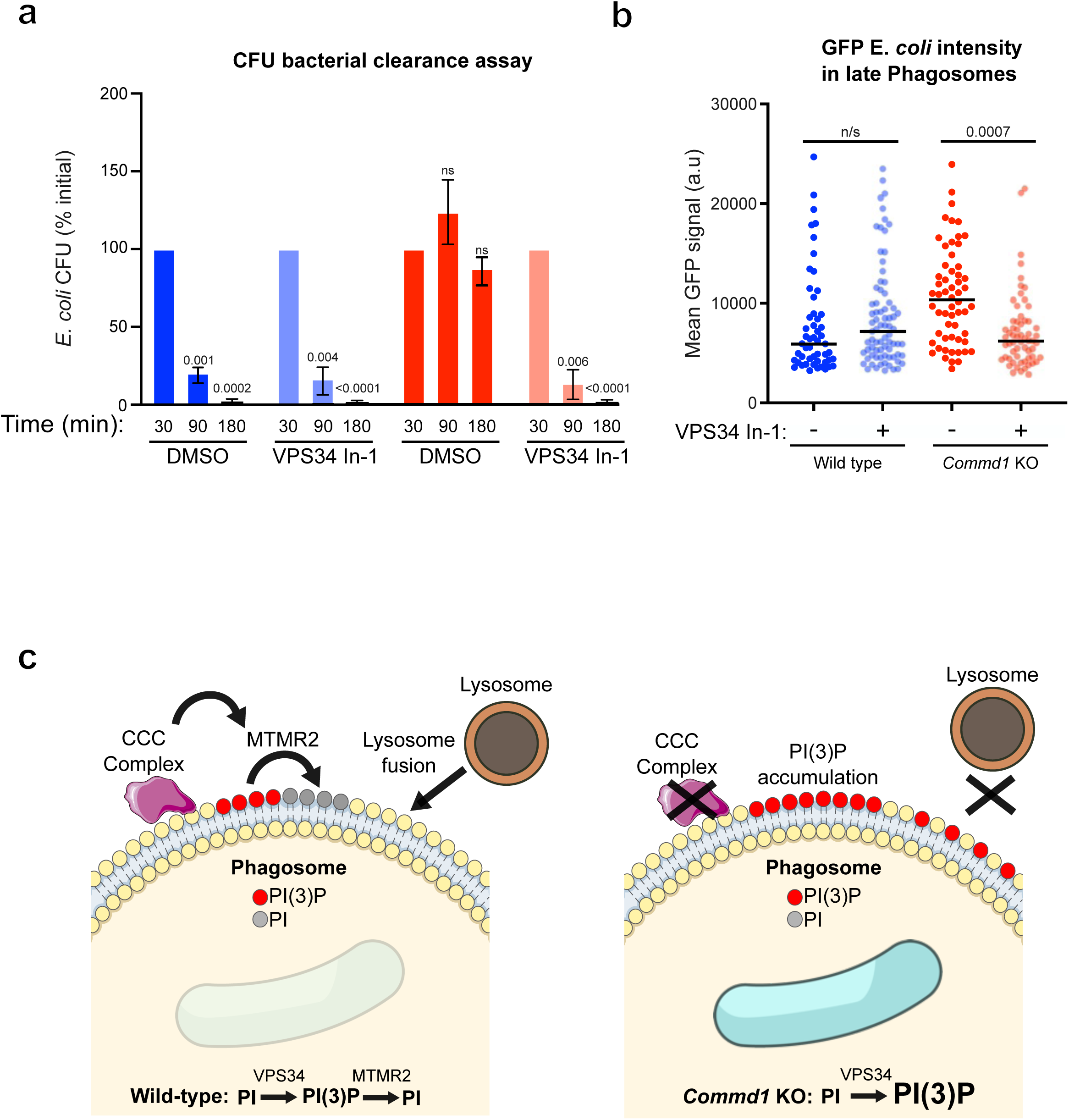
Restoration of PI(3)P homeostasis rescues bactericidal activity in *Commd1*-deficient macrophages. **(a)** Colony-forming unit (CFU) assay comparing bacterial clearance in wild-type and *Commd1* KO BMDMs treated with DMSO or VPS34-IN-1. Data are representative of two independent experiments from BMDMs isolated from three mice per genotype. (**b**) Quantification of GFP fluorescence intensity from three-dimensionally reconstructed phagocytosed GFP-expressing *E. coli* in live-cell imaging experiments performed in wild-type and *Commd1* KO BMDMs treated with DMSO or VPS34-IN-1 at late time points (10-min pulse, 90-min chase). Data are representative of two independent experiments from BMDMs isolated from three mice per genotype. (**c**) Model summarizing the proposed role of the CCC complex in regulating phagosomal PI(3)P turnover, phagosome maturation, and bactericidal activity. Two-tailed Student’s *t*-tests were used for pairwise comparisons in (**a**). One-way ANOVA was used for multiple-group comparisons in (**b**).

## Discussion

This study provides the first detailed mechanistic examination of the contribution of the CCC complex to phagosome maturation. We demonstrate that CCC, but not Retriever, is essential phagosome-lysosome fusion and ER-phagosome contact sites, which ultimately affect *E. coli* elimination. Critically, we demonstrate that the phenotype is mediated by reduced PI(3)P turnover secondary to defective recruitment of the lipid phosphatase, Mtmr2, and rebalancing PI(3)P levels on phagosome membranes is sufficient to restore the bactericidal activity of CCC deficient phagosomes.

The dispensable role of the Retriever complex in this context is particularly noteworthy, standing in stark contrast to the impact of CCC complex deficiency in phagosomal maturation. While CCC and Retriever were identified in separate studies, their strong physical association and cellular colocalization have promoted the view that these complexes predominantly exist as one larger protein assembly, referred to as Commander (Healy et al., 2023; Laulumaa et al., 2024; Mallam and Marcotte, 2017). Based on the distinct consequences observed when subunits of either complex are deleted in macrophages, our studies demonstrate that rather than acting as one functional unit, CCC and Retriever are separable functional units in the context of phagosome maturation.

Structural studies have uncovered that CCDC22 and CCDC93 heterodimerize to form an extended scaffold which on one end interacts with Retriever, and on the other end, interacts with a ring-like structure formed by all 10 COMMD proteins (Boesch et al., 2024; Healy et al., 2023; Laulumaa et al., 2024). Interestingly, while CCC subunits show strong colocalization on endosomes, confocal microscopy imaging of phagosomes demonstrates a different picture. On these larger structures, components of the CCC complex are not perfectly overlapping, with Ccdc22 and Ccdc93 displaying a more diffuse distribution throughout the phagosomal membrane, whereas Commd1 localizes to distinct focal areas that appear as puncta over that membrane. Consistent with this dissociation of the localization of Ccdc22, Ccdc93, and Ccommd proteins, our proteomic results indicate that Commd proteins have preferential enrichment in early phagosomes, whereas Ccdc22 and Ccdc93 have near equal abundance throughout phagosomal maturation. Thus, we speculate that the CCDC22/CCDC93 scaffold is recruited separately from the COMMD ring on phagosomal membranes, and eventually these components encounter themselves on specific membranes subdomains, resulting in the full assembly of the CCC complex.

One key observation in this study is the centrality of the PI(3)P accumulation caused by CCC deficiency to the functional impairment seen in *Commd1* deficient macrophages **(Figure 8).** Mechanistically, out studies point to reduced phagosomal recruitment of Mtmr2 as the proximate cause. These observations align well with a similar imbalance of PI(3)P on early endosomes previously found in CCC deficient HeLa cells (Singla et al., 2019), also linked to deficient MTMR2 recruitment (Franklin et al., 2013; Franklin et al., 2011; Singla et al., 2019). Rebalancing PI(3)P levels through partial VPS34 inhibition was sufficient to restore bactericidal activity in *Commd1* deficient macrophages. Interestingly, while we show that excess PI(3)P is detrimental to phagosome maturation, others have observed that severe PI(3)P deficiency due to loss of *VPS34*, has an equivalent effect of drastically reducing bactericidal activity (He et al., 2025; Vergne et al., 2005). Thus, precise PI(3)P regulation is necessary for proper phagosome maturation and fusion with lysosomes in macrophages. In addition, others have shown that although phagosome maturation is accompanied by a switch from PI(3)P to PI(4)P preponderance, both PI(3)P and PI(4)P are required to accomplish tethering and fusion of phagosomes and lysosomes (Jeschke and Haas, 2018). Although we could not accurately assess PI(4)P in late phagosomes due to poor signal-to-noise ratio of the PI(4)P biosensor, we postulate that the inability of CCC deficient macrophages to dephosphorylate PI(3)P may result in an imbalance between these critical lipids, preventing the normal steps of tethering and fusion of lysosomes. Additional evidence supporting this hypothesis comes from the observed reduction in ER-phagosomes contact sites noted in *Commd1* KO late phagosomes. Notably, multiple PI(4)P-binding factors, such as members of the OSBP family, VPS13C, and SAC1, are important in this process. The OSBP family, in particular, comprises lipid transfer proteins that have previously been implicated in regulating phagosomal PI(4)P levels (Lim et al., 2019).

Defects in phosphoinositide homeostasis have been shown to alter vesicle morphology and distribution in the cytoplasm, as has been described for CCC deficiency (Singla et al., 2019) as well as PIKfyve deficiency (Choy et al., 2018). Consistent with this idea, we observed that PI(3)P-positive vesicles in Commd1 KO cells were enlarged relative to those in wild-type cells (**Extended Data Figure 6**). Similarly, late *Commd1* KO phagosomes were approximately two-fold larger than their wild-type counterparts (**Figure 5f**). Together, these observations indicate that altered PI(3)P turnover in CCC deficient macrophages disrupts the recruitment of ER–phagosome contact site proteins involved in lipid transport and exchange. Such defects could impair normal membrane remodeling during phagosome maturation, ultimately contributing to phagosome enlargement. Although further work will be required to directly test this model, the potential interplay between the CCC complex, phosphoinositide metabolism, and ER-mediated lipid transfer represents an intriguing area for future investigation.

Based on these observations, one prediction would be that disabling the CCC complex assembly, its phagosomal recruitment, or the recruitment and activity of MTMR2, may provide pathogens with a growth advantage in macrophages. To date, effector proteins targeting CCC subunits remain to be identified, and the closest mechanism known is the ability of *Burkholderia cenocepacia* to target the WASH complex (Walpole et al., 2020), which is required for CCC recruitment to endosomes (Phillips-Krawczak et al., 2015). By extension, another prediction is that pathogens may benefit from manipulating PI(3)P levels to prevent phagosome-lysosome fusion and promote microbial survival. Indeed, organisms that block phagosome-lysosome fusion, such as *M. tuberculosis* or *L. pneumophila,* express pathogenicity effector proteins that function as PI(3)P phosphatases (Deretic, 2008; Swart and Hilbi, 2020; Toulabi et al., 2013; Vergne et al., 2003; Vergne et al., 2005). While CCC mutations in humans cause developmental defects, so called Ritscher-Schinzel or 3C syndrome (Kato et al., 2025; Singla et al., 2025; Starokadomskyy et al., 2013), immune defects have yet to be reported; nonetheless, the total number of known cases, with less than 30 patients reported with severe mutations in CCC subunits (Gjerulfsen et al., 2021; Kato et al., 2025; Kolanczyk et al., 2014; Singla et al., 2025; Voineagu et al., 2012). In the case of *MTMR2* mutations, severe progressive peripheral polyneuropathy known as Charcot-Marie-Tooth 4B1 syndrome (Pareyson et al., 2019) has been reported, but again with fewer than 60 patients reported, no immune defects have been noted (Du et al., 2024; Guimaraes-Costa et al., 2020; Halperin et al., 2020; Pareyson et al., 2019). Thus, future characterization of immune function in these patients, particularly for macrophage defects, is warranted.

In conclusion, this study establishes a unique role for the CCC complex in phagosome maturation through PI(3)P regulation and opens the door for future investigation of potential microbial effectors targeting this pathway as well as further analysis of how this pathway may be altered in inborn errors of immunity.

## Methods

### Regents

Antibodies used in this study are described in **Supplementary Table 3**. Other reagents used include green-fluorescent carboxylate-modified polystyrene latex beads (Sigma Aldrich, L4655), magnetic beads (Polysciences, 25029-5), Tamoxifen (Sigma Aldrich, T5648), and the VPS34 inhibitor, VPS34 IN-1 (Cayman Chemical Company, Cat# 17392). GFP-expressing *E. coli* were obtained from ATCC (Cat# 25922GFP).

### Mice

*Commd1^fl/fl^* mice have been previously reported (Li et al., 2014; Vonk et al., 2011). Mice carrying conditional alleles in the *Vps35l* or *Vps26c* genes were generated in the transgenic core facility at UT Southwestern, using the easi-CRISPR approach (Quadros et al., 2017). Details of the LoxP locations in our targeting strategy are provided in **Extended Data Figure 4**. Guide RNA sequences are provided in **Supplementary Table 4**. In the case of Vps26c, an in-frame HA tag coding sequence was added at the 5’ end of the coding sequence given the poor sensitivity of available anti-Vps26c antibodies. Targeted regions in founders were fully sequenced to ensure that the alleles corresponded to the desired results. Thereafter, animals were backcrossed to C57BL6 mice for at least 3 generations. Mutant mice were mated to *Lys2^Cre^* mice (Clausen et al., 1999) to generate constitutive myeloid deficiency of the CCC complex, or to *Ubc^CreERT2^* mice (Ruzankina et al., 2007), where post-natal IP injection with 100μL tamoxifen (20 mg/mL in corn oil) every day for 5 days was administered to induce gene deletion. Bone marrow was harvested 10 days following the completion of the tamoxifen administration.

### Cell culture

L929 cells were grown in 90% DMEM, 10% FBS, supplemented with glutamax (1 g/dL) for 10 days at a ratio of 2.8 mL of media per square centimeter of area of confluent L929 cells. Afterwards, conditioned media was harvested and stored at -80°C. BMDMs were generated by harvesting bone marrow from mouse legbones, which was then placed in BMDM growth media consisting of 45% DMEM, 20% FBS, and 35% L929 conditioned media. Bone marrow cells were allowed to differentiate for 7 days. During this time, BMDM would attach to tissue culture plates and acquire typical morphology. Thereafter, BMDMs were split for experiments using a cell scraper to harvest from plates.

### Bacterial Clearance assays

For plate reader assays, GFP expressing *E. coli* were grown in LB broth overnight at 37°C. From this overnight culture, 300mL was added to 10mL LB broth and incubated for 1.5 h at 37°C to allow for the bacteria to reach their exponential growth phase. From this new culture, 1mL was centrifuged at 4000xg for 2 min, and the pelleted bacteria were washed three times with PBS. The pellet was finally resuspended to achieve an OD600 of 1, corresponding to ∼8x10^8^ bacteria/mL. Bacteria were then added to a 96 well plate containing either wild-type or specified KO BMDMs, at an MOI of 20. The plates were then centrifuged using a swing bucket rotor for 5 minutes at 1000xg to bring the bacteria in contact with the attached macrophages. Afterwards, bacteria that were not phagocytosed by BMDMs were aspirated and the wells were rinsed with PBS three times. After rinsing, fresh media supplemented with gentamycin (20mg/mL) was added to the wells and the plate was placed on a BioTek Synergy H1 microplate reader at 37°C. At this point, the plate reader was used to measure GFP signals at specified time-points. For the CFU assays, bacteria were prepared similarly but were instead added to wells on a 24-well plate containing either wild-type or KO BMDMs, at an MOI of 20. Centrifugation to promote rapid contact of bacteria with macrophages was performed as outlined above, and cells were incubated for 30 minutes at 37°C before excess bacteria were removed by aspiration and PBS rinsing (three times), followed by addition of fresh BMDM media containing gentamycin (20μg/mL). At the indicated timepoints, the media was removed, and BMDMs were lysed by adding 0.05% Triton X in PBS and allowed to sit at room temperature for 15 minutes. At this point, lysates were harvested and plated in serial 10-fold dilutions on agar plates to allow bacteria to grow overnight. The next morning, colonies were counted manually. For experiments involving VPS34 IN-1, cells were treated with 500 nM VPS34 IN-1 or an equivalent volume of DMSO for 30 minutes prior to and throughout the duration of the experiment.

### Phagosome Isolation

In some experiments, sucrose gradient centrifugation was used for phagosome isolation, following a previously reported protocol (Jeschke et al., 2015). In short, 1μm green-fluorescent carboxylate-modified polystyrene latex beads were added at 6x10^8^ beads/mL to a 15-cm plate of BMDMs at 95% confluency for 10 minutes. Subsequently, BMDMs were either harvested or were rinsed with PBS three times, given fresh media, and returned to culture conditions for another 80 minutes. BMDM harvesting was performed by placing the culture plates on ice, removing the growth media and adding ice-cold PBS, followed by scraping using a cell scraper. Harvested BMDMs were washed three times by centrifugation at 160xg for 7 minutes at 4°C, first with PBS, next with PBS + 5mM EDTA, and last with HB buffer (20 mM HEPES/KOH (pH 7.2), 8.6% (wt/vol) sucrose, 0.5 mM EGTA). After washing, cells were mixed with HB buffer + protease inhibitors and homogenized with a 2mL Dounce homogenizer (approximately 20-25 strokes). Homogenized lysate was centrifuged at 1000xg for 5 minutes at 4°C. The supernatant was harvested, adjusted to 32% sucrose using HB buffer and transferred to a high-speed centrifuge tube; 25% and 8.6% sucrose in HB buffer were sequentially layered on top of the lysate before centrifugation at 42,000xg for 30 minutes at 4°C. The phagosome fraction, visible between the 8.6% and 25% sucrose interface, was harvested using a pipet reaching into the 8.6-25% sucrose interface and slowly collecting the phagosomes until no more green phagosomes where observed. The harvested phagosome fraction was diluted 6-fold with PBS supplemented with protease inhibitors. This phagosome fraction was centrifuged at 3000xg for 30 minutes at 4°C in a glass tube to pellet phagosomes while avoiding excessive adherence to the tube wall. Phagosome pellets were then resuspended with LDS gel loading buffer (Invitrogen, NP0008), boiled, and loaded on polyacrylamide gels for proteomic or immunoblot analyses.

For other experiments, magnetic beads were used for phagosome isolation (Liu and Liu, 2025). BMDMs were incubated with 1μm magnetic beads at 6x10^8^ beads/mL for 10 minutes. Subsequently, BMDMs were rinsed with PBS three times and either harvested on ice with a cell scraper or given fresh media and returned to culture conditions for another 80 minutes before harvesting. Harvested BMDMs were resuspended in hypotonic HB buffer (20 mM HEPES/KOH (pH 7.2), 8.6% (wt/vol) sucrose, 0.5 mM EGTA) and homogenized with a 2mL Dounce homogenizer as described above. This homogenized lysate was then centrifuged at 4°C at 200xg for 5 minutes to precipitate nuclei and other large debris; the phagosomes were harvested from the cytosolic fraction by applying a magnetic rack on ice, followed by 3 washes with ice cold PBS. For flow cytometry staining, isolated phagosomes were incubated with primary antibody diluted in PBS supplemented with 5% goat serum for 1 hour on ice, followed by rinsing in cold PBS + 5% goat serum and pelleting using the magnetic rack. The rinsing step was repeated 3 times. Phagosomes were then incubated with fluorescent secondary antibodies diluted in PBS + 5% goat serum for 30 minutes on ice. Three washes using PBS + 5% goat serum were performed before proceeding to flow cytometry.

### Proteomic analysis

Samples were digested overnight with trypsin (Pierce) following reduction and alkylation with DTT and iodoacetamide (Sigma–Aldrich). Following solid-phase extraction cleanup with an Oasis HLB µelution plate (Waters), the resulting peptides were reconstituted in 10 mL of 2% (v/v) acetonitrile (ACN) and 0.1% trifluoroacetic acid in water. Of this sample, 5 mL was injected onto either an Orbitrap Fusion Lumos or Orbitrap Eclipse mass spectrometer (Thermo) coupled to an Ultimate 3000 RSLC-Nano liquid chromatography systems (Thermo). Samples were injected onto a 75 μm i.d., 75-cm long EasySpray column (Thermo): mobile Phase A contained 2% (v/v) ACN and 0.1% formic acid in water, and Mobile Phase B contained 80% (v/v) ACN, 10% (v/v) trifluoroethanol, and 0.1% formic acid in water. Peptides were eluted with a gradient from 0-28% mobile phase B over 90 min. The mass spectrometer was operated in positive ion mode with a source voltage of 2.5 kV and an ion transfer tube temperature of 300°C. MS scans were acquired at 120,000 resolution in the Orbitrap and up to 10 MS/MS spectra were obtained in the Orbitrap for each full spectrum acquired using higher-energy collisional dissociation (HCD) for ions with charges 2-7. Dynamic exclusion was set for 25 seconds after an ion was selected for fragmentation.

Raw MS data files were analyzed using Proteome Discoverer v.3.0 (Thermo), with peptide identification performed using Sequest HT searching against the mouse reviewed protein database from UniProt. Fragment and precursor tolerances of 10 ppm and 0.6 Da were specified, and three missed cleavages were allowed. Carbamidomethylation of Cys was set as a fixed modification, with oxidation of Met set as a variable modification. The false-discovery rate (FDR) cutoff was 1% for all peptides. The sum of peak intensities for all peptides matched to a protein were used as protein abundance values. Normalization was performed such that the sums of the abundance values were equal for all samples in the comparison. These normalized values were input into Argonaut (Brademan et al., 2020) (version 2.0) for further analysis, with missing values imputed based on a left-censored missing value algorithm.

Replicates were compared using a PCA plot and outliers were removed before analysis. Analysis of proteomics hits for pathway identification was done using Enrichr (Chen et al., 2013), focusing on GO Cellular Component outputs. Volcano plots were prepared using normalized abundance values and FDR corrected p values and plotted using VolcaNoseR (Goedhart and Luijsterburg, 2020). Clustering of protein abundance results and heat maps generation was accomplished using Heatmapper (Babicki et al., 2016) (version 2).

### Immunostaining

A protocol adapted from a prior publication was used (Singla et al., 2019). Cells were fixed in ice-cold fixative (4% paraformaldehyde in PBS) and incubated for 18 min at room temperature in the dark, followed by permeabilization for 3 min with 0.15% Surfact-Amps X-100 (Thermofisher Scientific) in PBS. Samples were then washed three times and incubated overnight at 4°C in a humidified chamber with primary antibodies in immunofluorescence (IF) buffer (consisting of Tris-buffered saline plus 5% Goat serum). After three washes in PBS, the samples were incubated with secondary antibodies (1:500 dilution in IF buffer) for 1 h at room temperature or overnight at 4°C in a humidified chamber; afterwards four PBS washes were done.

### PI(3)P probe staining

The 6xHis-SNAP-PI(3)P probe was expressed as described previously (Maib et al., 2024), using a plasmid deposited in Addgene by Hannes Maib & David Murray (Addgene plasmid # 211508). BL21 Gold DE3 *E. coli* cells (Agilent) were transformed with this plasmid, and the expression of recombinant proteins was induced with 1mM IPTG at 18°C for 16 hr. Cells were lysed by sonication. Recombinant proteins were purified using Ni-NTA beads (Qiagen), followed by cation exchange chromatography using a Source 15S column (Cytiva) at pH 7.0, and then by size exclusion chromatography (SEC) using a Superdex200 column (Cytiva) into 100 mM NaCl, 20 mM PIPES pH 6.8, 2 mM MgCl_2_, 5% (w/v) glycerol, and 1mM DTT. The protein was fluorescently labeled using SNAP-Surface® Alexa Fluor® 647 (NEB) following manufacturer recommendations. Briefly, reactions were set up using 5 µM protein, 10 µM SNAP-Surface® Alexa Fluor® 647 dye, and 1 mM DTT in the same buffer as the SEC was performed. The reaction was incubated for 2 hours at room temperature, at which point the protein was desalted using a HiTrap Desalting column (Cytiva) into 100 mM NaCl, 20 mM PIPES pH 6.8, 2 mM MgCl_2_, 5% (w/v) glycerol, and 1mM DTT, and concentrated for use. Protein and dye concentrations were determined using a NanoDrop, with an extinction coefficient of 255,000 M^-1^ cm^-1^ at 650nm used for the dye.

Staining for PI(3)P was performed as described (Maib et al., 2024). In short, BMDMs were plated on imaging plates (Cellvis) to reach ∼75% confluency. GFP-expressing *E. coli* were prepared as described in the bacterial clearance assay and added to the plated BMDMs at an MOI of 10 for the indicated pulse/chase times. This was followed by media aspiration and fixation using 4% PFA in PBS at room temperature for 20 minutes. BMDMs were then washed three times with ice cold PIPES buffer (20mM of Pipes, pH^+^ 6.8, 137mM NaCl, 2.7mM KCl), and then incubated for 45–60 min on ice with PIPES buffer containing 5% BSA + 0.5% (vol/wt) saponin with the addition of fluorescently labeled PI(3)P probe to a final concentration of 500 nM. This achieved blocking, permeabilization and staining in one step. Samples washed 3x with PBS before imaging. For experiments involving VPS34 IN-1, cells were treated with 500 nM VPS34 IN-1 or an equivalent volume of DMSO for 30 minutes prior to and throughout the duration of the experiment.

### Confocal microscopy

Confocal imaging of fixed cells was performed at room temperature with a CFI Apo 100× objective (1.49 NA) and a custom spinning-disk confocal-TIRF imaging system built around an Eclipse Ti microscope (Nikon) with an ORCA-Fusion BT digital CMOS camera (Hamamatsu Photonics). The microscope was controlled by NIS-Elements (version 5.42.04, Nikon Instruments Inc). Excitation of fluorescence was achieved using a LU-NV laser unit. For live cell imaging, a humidified, CO_2_, and temperature (37°C) controlled chamber (Okolab USA Inc) was placed over the sample handler of the microscope. Fixed cells were imaged in PBS while live cell imaging was done in BMDM growth media.

### Image analysis

Quantification of fluorescence intensity on the phagosome membrane (Tmem192 and PI(3)P) was done using Fiji (version 2.16.0). Using z-stack images of macrophages, a slice best representing *E. coli* inside each macrophage was chosen. Signal thresholding was done for GFP expressing *E. coli* from max intensity to the predetermined value of 1900 pixel intensity. This value was chosen as it was above cell background and allowed for low intensity bacteria to be included in analysis. Bacteria outside of cells were manually excluded from analysis. At this point, a 0.5 μm band was created around single bacteria and the signal of the marker of interest (Tmem192 or PI(3)P) was measured in this band. Background signal was subtracted using the signal intensity between cells for the channel of interest.

For specific experiments, 3D reconstruction and image analysis was performed using Imaris (version 10.2), in which a surface was created for either *E. coli* or TMEM192. Signal thresholding was done for *E. coli* from max intensity to the predetermined value of 1900 pixel intensity. For TMEM192 background subtraction for “diameter of largest sphere which fits into object” was set to 0.5 µm, signal thresholding was automatically set, and “split touching objects” seed diameter was set to 0.5 µm. For cell size a mask was drawn around the cell using phalloidin as a cell boundary marker.

### Electron microscopy

Cells were fixed on MatTek dishes with 2.5% (v/v) glutaraldehyde in 0.1M sodium cacodylate buffer. After five rinses in 0.1 M sodium cacodylate buffer, cells were post-fixed in 1% osmium tetroxide and 1.5% K_3_[Fe(CN)_6_] in 0.1 M sodium cacodylate buffer for 1 hour at 4°C. Cells were rinsed with water and en bloc stained with 0.5% aqueous uranyl acetate in 25% methanol overnight at 4°C. After five rinses with water, specimens were stained with 0.02M lead nitrate in 0.03M L-aspartate acid in a 60°C oven for 30 minutes. After 5 water washes, samples were dehydrated with increasing concentration of ethanol, infiltrated with Embed-812 resin and polymerized in a 60°C oven overnight. Embed-812 discs were removed from MatTek plastic housing by submerging the dish in liquid nitrogen. Pieces of the disc were glued to blanks with super glue. Blocks were sectioned with a diamond knife (Diatome) on a Leica Ultracut UCT (7) ultramicrotome (Leica Microsystems), collected onto copper grids, and post stained if needed with 2% Uranyl acetate in water and lead citrate. Images were acquired on a JEM-1400 Plus transmission electron microscope equipped with a LaB6 source operated at 120 kV using an AMT-BioSprint 16M CCD camera.

### Immunoblotting

Immunoblotting experiments were performed by resolving the protein samples using 4–12% gradient Novex Bis-Tris gels (Invitrogen), transferred to nitrocellulose membranes and blocked with 5% milk solution in Tris-buffered Saline buffer containing 0.05% Tween-20. Subsequently, membranes were incubated with primary antibodies, followed by incubation with HRP-conjugated secondary antibodies.

### Flow cytometry

Samples were acquired on a 5-Laser Aurora (Cytek Bio) flow cytometer (using SpectroFlo software version 3.3.0) calibrated using SpectroFlo QC. Calibration of the cytometer before acquisition was performed with Submicron Bead Calibration Kit (Bangs Laboratories). Data was analyzed using Flowjo software (version v10.10). During analysis, phagosomes were gated to only include phagosomes containing one bead. Median of the 647nm channel (comp-R2) was used for comparing WT and KO phagosomes.

### Statistical analysis

Prism (version 10.6.1) was used for plot generation and statistical analyses. When comparing two groups with small sample sizes, we used Student’s *t*-test; for multigroup analyses, we used one way ANOVA. For datasets with large number of variables, false discovery rate correction (FDR) using the Benjamini-Hochberg method was used. The software packages used for these analyses are described in the proteomics section above.

## Acknowledgements

We are grateful to Drs. Mike Henne, Hans-Christian Reinecker, Joseph Albanesi, Marcel Mettlen, Josephine Ni and Jungyeon Kim for their helpful input during the development of this project. We are also grateful for the following UT Southwestern core facilities and their staff: Proteomics core, Flow cytometry Core, Quantitative Light Microscopy Core and EM core (supported by NIH S10OD021685 and P30CA142543 grants). The work was supported by NIH grants (R01DK107733, R01HL179813 to EB; R01GM144479, R01NS137605 to JL, R01GM128786, R01HL179813 to BC, R01AI083359 to NA), the Welch Foundation (grant no. I-1704 to NA) and by institutional resources (Pollock Family Center for Research in Inflammatory Bowel Disease to EB). JL is a UT Southwestern Sowell Family Scholar in Medical Research.

## Author Contributions

EB and JL conceived the study and supervised the project. Experimental design was led by EB, JL and also contributed by NA, BC and HB. HB conducted the bulk of the experiments. Some specific experiments were conducted by WB, DAK, RP, AS, QL, LS, W-RL, JY, ST, BC. HB analyzed the data and prepared the figures. EB, JL and HB interpreted the results and wrote the manuscript.

## Data Availability

Proteomics results presented here have been deposited in the Massive repository (accession number MSV000099882). Source data are provided with this paper. All other data supporting the findings of this study are available from the corresponding author upon reasonable request.

## Supplementary Data

Supplementary Table 1: Proteomics data table comparing phagosomes from wild-type and Commd1 deficient macrophages (related to Figures 3, 6, and Extended Data Figure 5).

Supplementary Table 2: Proteomics data table comparing phagosomes from wild-type and *Vps35l* deficient macrophages (related to Figures 5, and Extended Data Figure 5).

Supplementary Table 3: Antibodies used in this study.

Supplementary Table 4: Guide RNA sequences used to generate *Vps35l* and *Vps26c* conditional mouse lines

**Extended Data Fig. 1:**
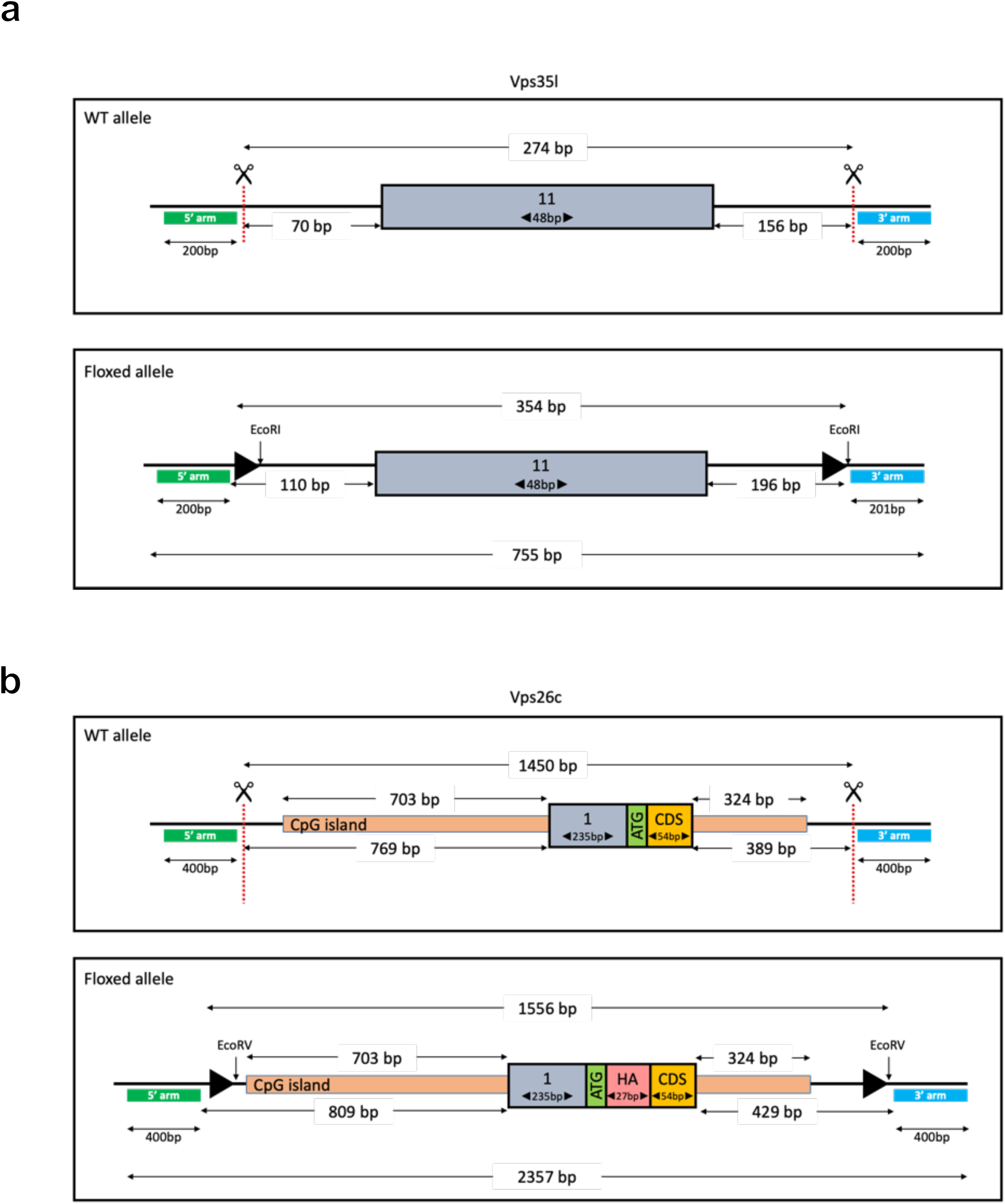
Generation of conditional Vps35l and Vps26c alleles. (**a**, **b**) Schematic representation of the targeting strategies used to generate the conditional *Vps35l* (**a**) and *Vps26c* (**b**) alleles.

**Extended Data Fig. 2:**
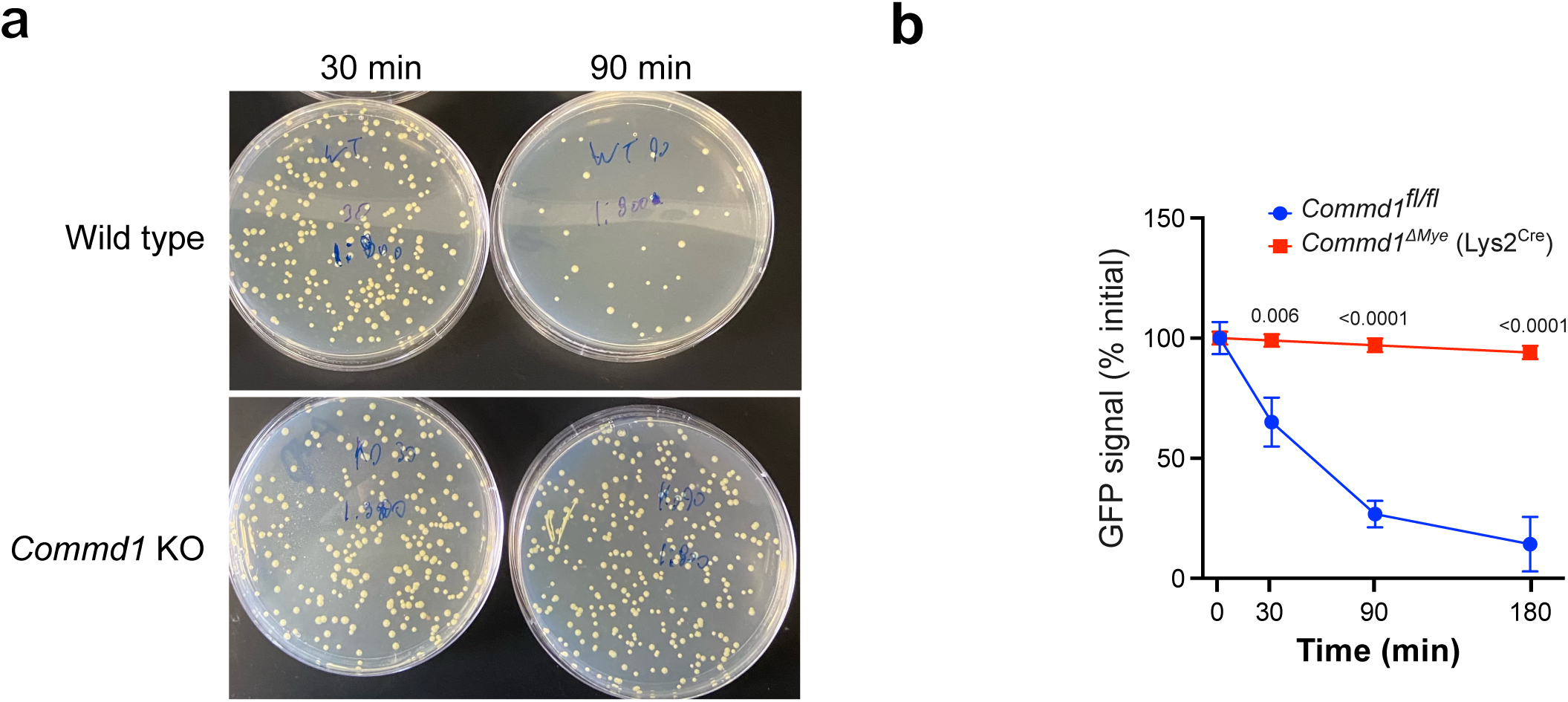
Impaired intracellular bacterial clearance in Commd1-deficient macrophages. (**a**) Representative images of agar plates used for colony-forming unit (CFU) assays. Plates from wild-type and Commd1-knockout BMDMs are shown at the indicated time points. (**b**) GFP fluorescence from phagocytosed GFP-expressing *E. coli* was monitored over time in live macrophages using a fluorescence plate reader. Wild-type and *Commd1* knockout BMDMs were compared. Data represents two independent experiments from BMDMs isolated from three mice per genotype. Two-tailed Student’s t-tests were used for pairwise comparisons between genotypes at each time point.

**Extended Data Fig. 3:**
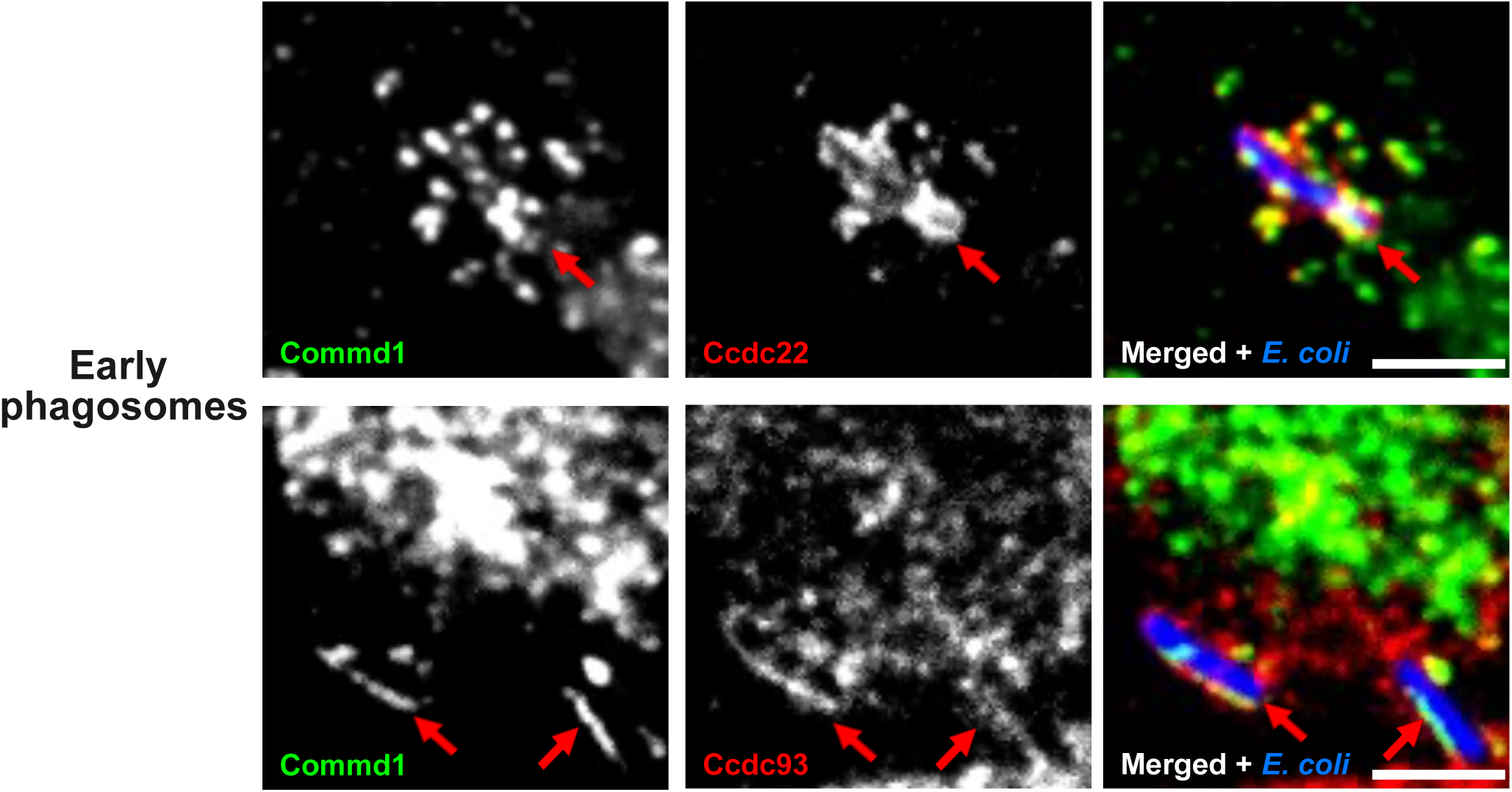
Localization of CCC complex components to *E. coli*-containing phagosomes. (**a**) Representative immunofluorescence images of early phagosomes (10 min) containing *E. coli* (blue). The upper panels show staining for Commd1 (green) and Ccdc22 (red). The lower panels show staining for Commd1 (green) and Ccdc93 (red). Scale bar, 2 μm.

**Extended Data Fig. 4:**
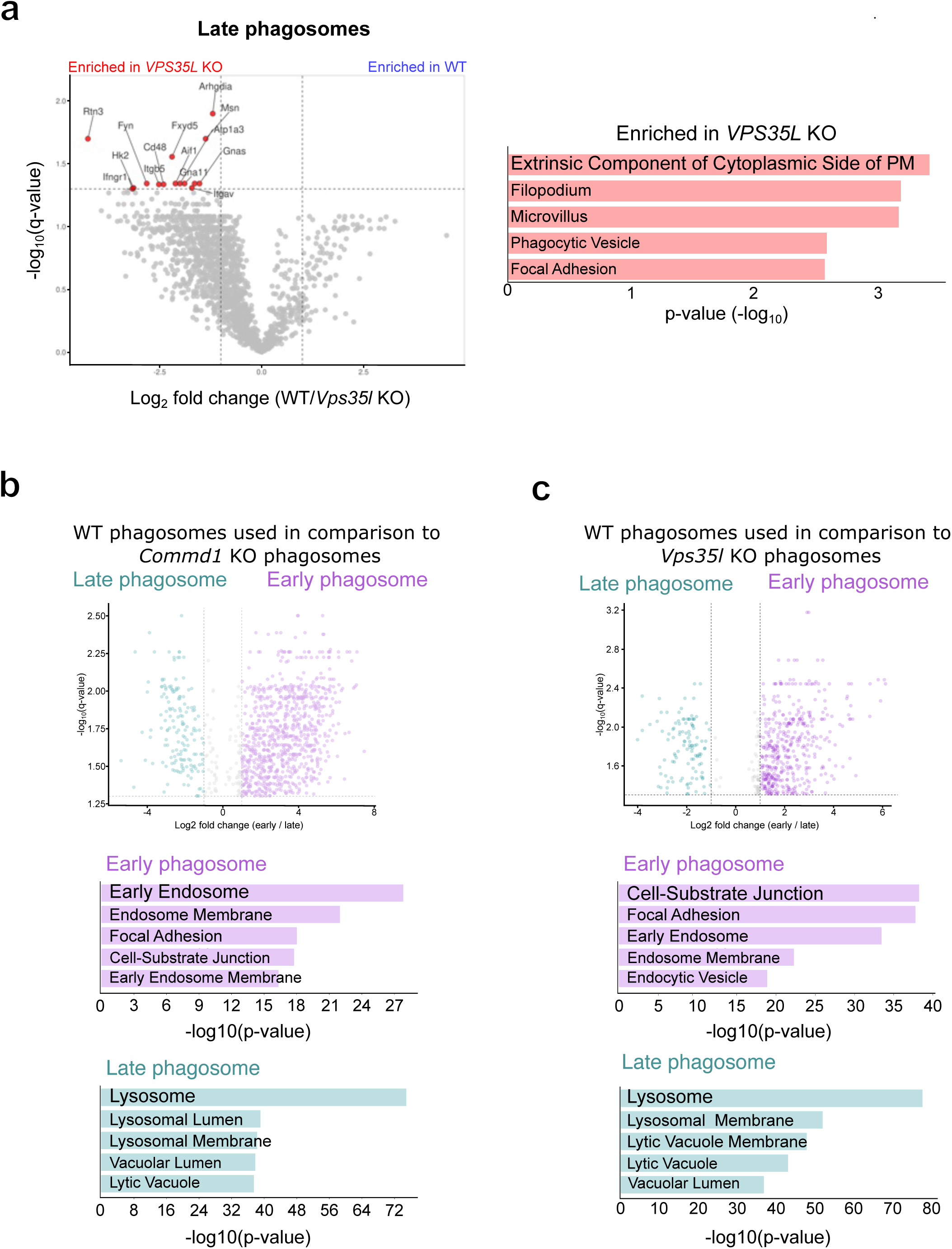
Proteomic analysis of phagosomes isolated from Vps35l-deficient macrophages. (**a**) Annotated volcano plot showing proteins significantly enriched in late phagosomes isolated from *Vps35l* knockout BMDMs relative to wild-type controls (top). Corresponding pathway enrichment analysis is shown below. (**b**) Volcano plot (top) and pathway enrichment analysis (bottom) comparing early and late phagosomes isolated from wild-type (*Commd1* fl/fl) macrophages. (**c**) Equivalent analysis comparing early and late phagosomes isolated from an independent wild-type control line (*Vps35l* fl/fl).

**Extended Data Fig. 5:**
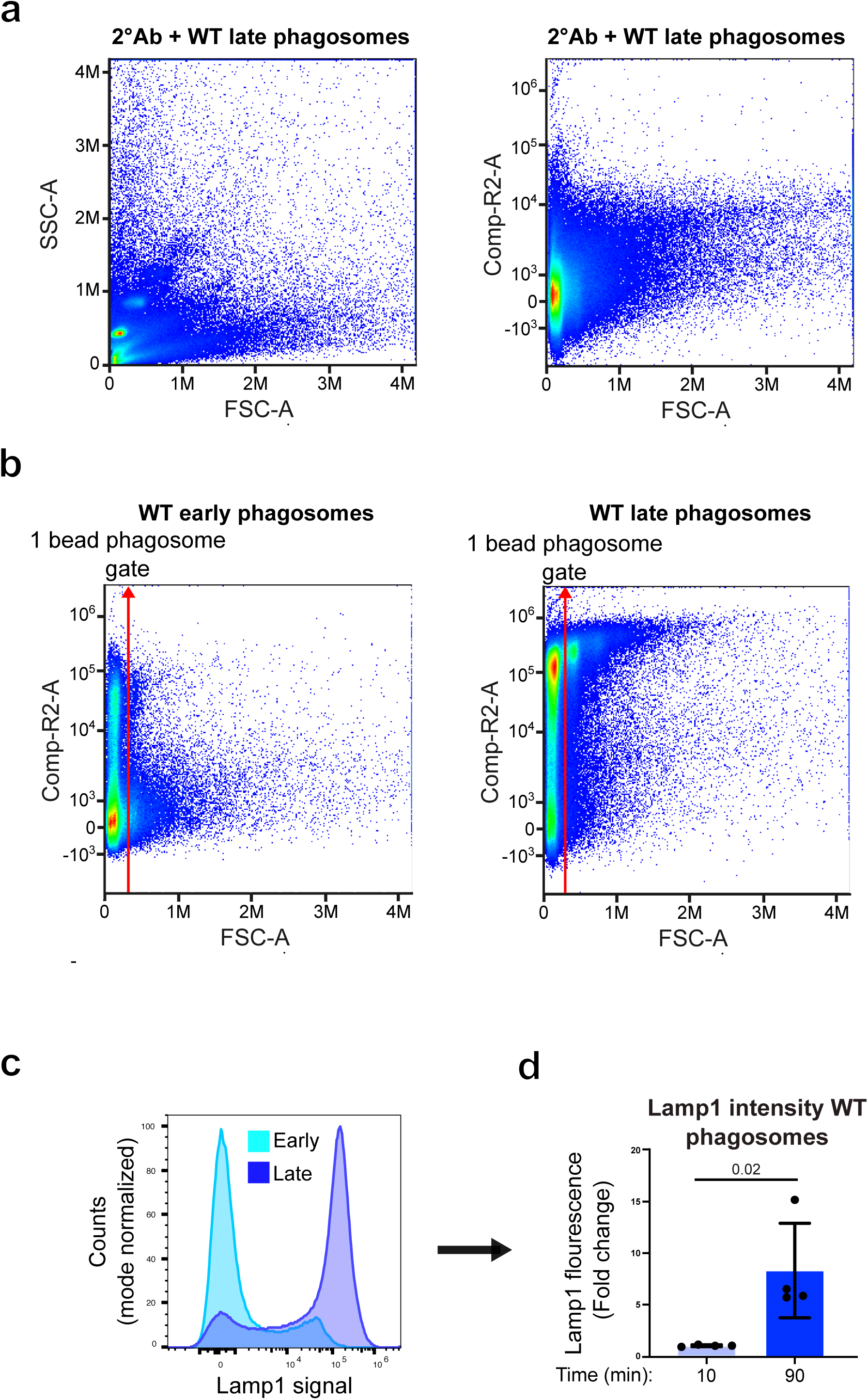
Flow-cytometric gating strategy for analysis of Lamp1-positive phagosomes. (**a**) Representative forward- and side-scatter plots of magnetic bead-containing phagosomes stained for Lamp1 (left) or stained with secondary antibody alone (right). (**b**) Gating strategy used to analyze Lamp1-positive phagosomes. Analysis was restricted to phagosomes containing a single magnetic bead. Representative plots of early (left) and late (right) wild-type phagosomes are shown. Student’s *t*-test was used for pairwise comparison between genotypes and timepoints at each time point in **(b).**

**Extended Data Fig. 6:**
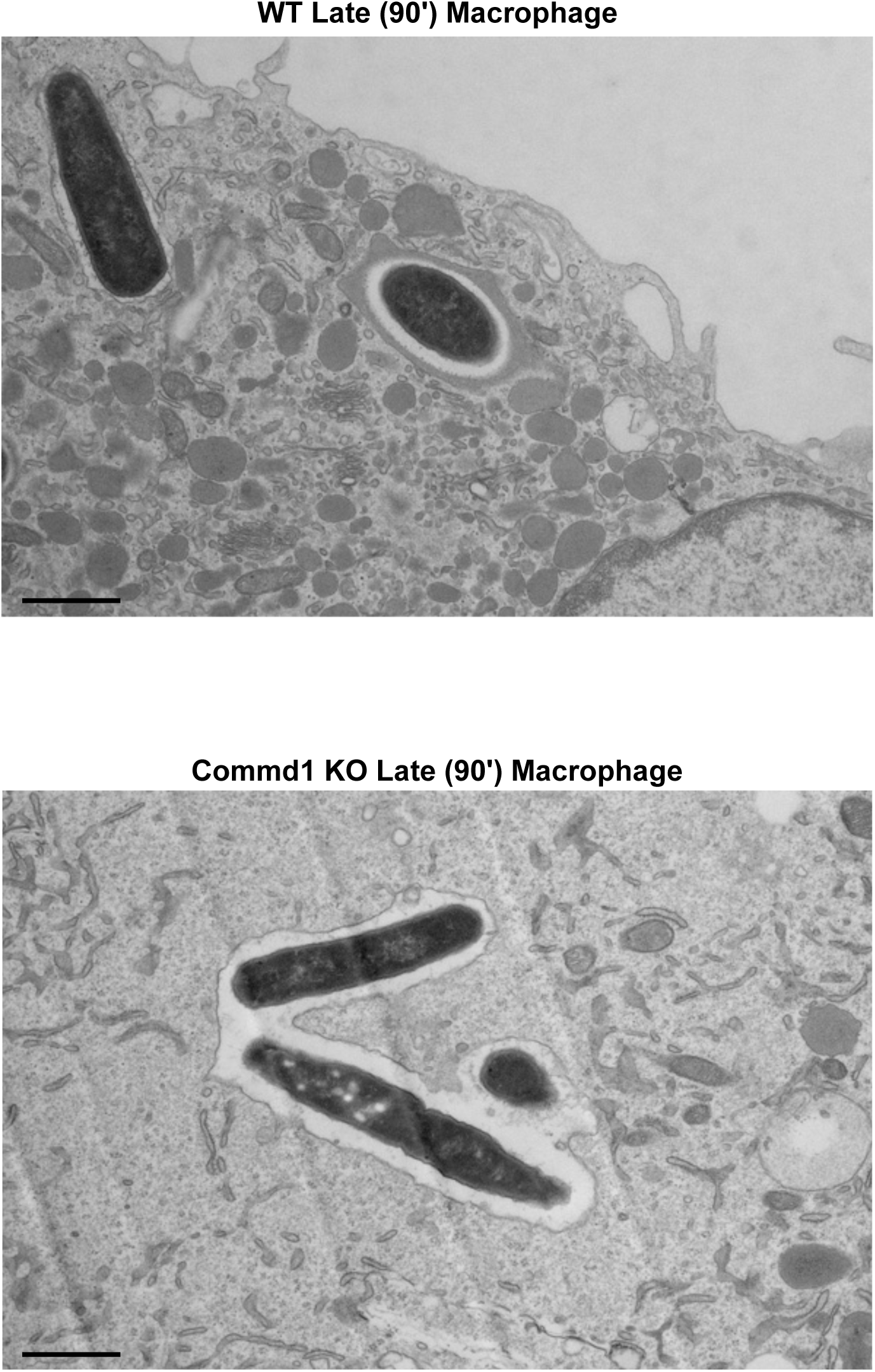
Lysosome proximity to late phagosomes. **(a)** Representative images of wild type and *Commd1* KO macrophages that have phagocytosed bacteria for 90 minutes.

**Extended Data Fig. 7:**
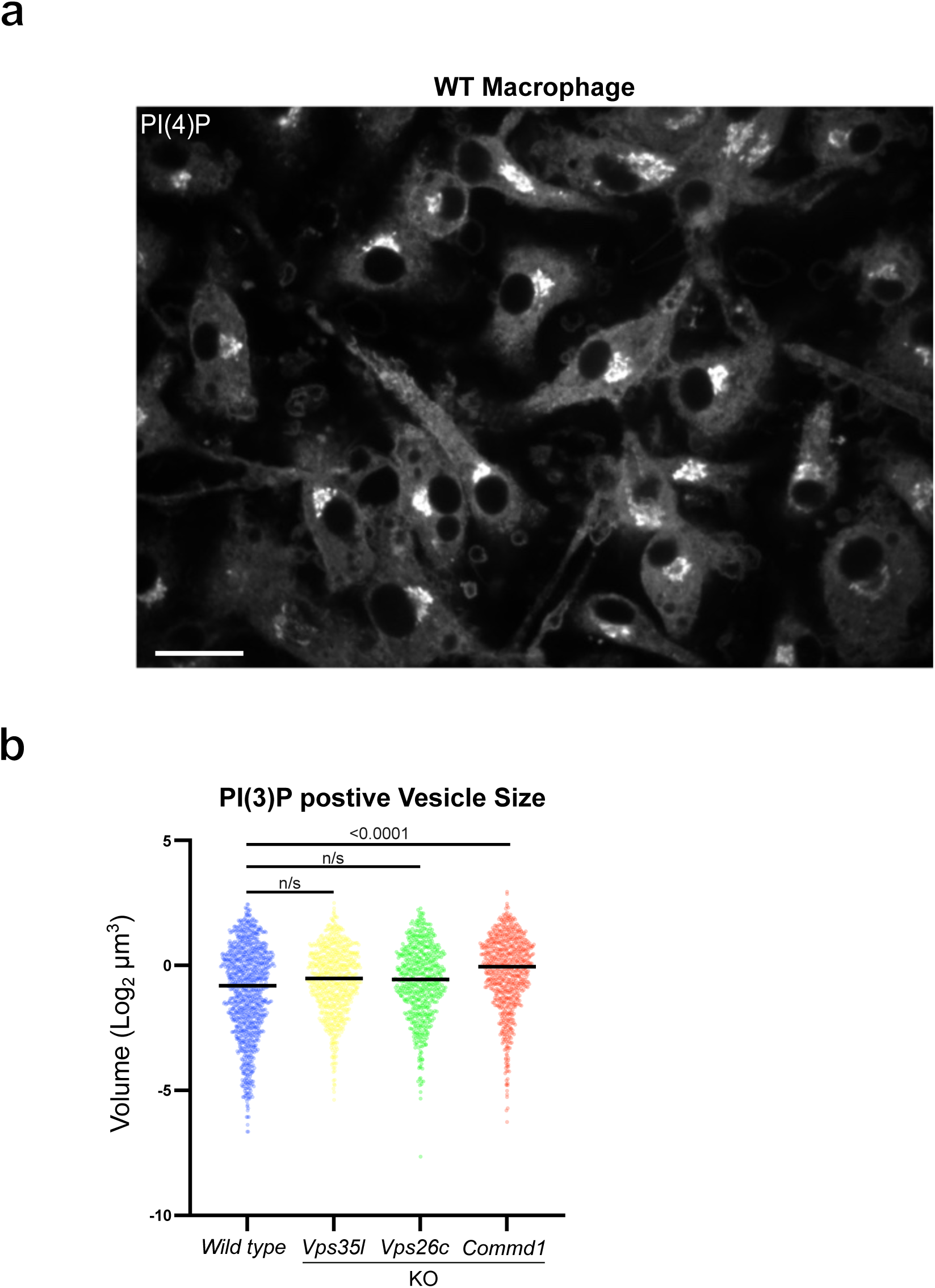
Increased PI(3)P-positive vesicle size in CCC-deficient macrophages. (**a**) Representative image of wild type macrophages stained with a PI(4)P binding probe. Scale bar 20 μm. (**b**) Quantification of the median volume of three-dimensionally reconstructed PI(3)P-positive vesicles in wild-type, *Commd1* knockout, *Vps35l* knockout and *Vps26c* knockout BMDMs. Data are representative of 20 cells from two independent experiments from BMDMs isolated from two mice per genotype. One-way ANOVA was used for multiple-group comparisons in (**b**).

