## Supplemental Table 4 for "Phosphoinositide regulation by the CCC complex promotes phagosome maturation and host defense"

**Supplementary Table 4:** Guide RNA sequences used to generate *Vps35l* and *Vps26c* conditional mouse lines

| Target | Strand | Sequence | PAM | Specificity | Efficiency |
|---|---|---|---|---|---|
| Vps26c | + | CAGTGGATACAAATGCAGCG | TGG | 76.6 | 74.6 |
| Vps26c | + | GAATAAAGTGTATCACGCCG | GGG | 92.9 | 68.2 |
| Vps35l | - | TCTGCCTGCCTCAAGCACCG | GGG | 69 | 71.7 |
| Vps35l | - | CAGTTTCCACACACTACGTG | GGG | 80.5 | 66.8 |
