## Supplemental Table 3 for "Phosphoinositide regulation by the CCC complex promotes phagosome maturation and host defense"

**Supplementary Table 3:** Antibodies used in this study.

| Target protein | Antibody source (clone) | Immunoblotting use | Immunostaining use | Validation reference |
|---|---|---|---|---|
| Rab5 | Cell signaling, 3547 (rabbit polyclonal) | 1:1000 dilution | n/a | Wenlu Ouyang, et al. 2025 |
| Rab7 | Cell signaling, 9367 (rabbit polyclonal) | 1:1000 dilution | n/a | Kaushal Asrani, et al. 2025 |
| MTMR2 | Santa Cruz, sc365185, (mouse monoclonal) | 1:1000 dilution | n/a | |
| Tmem192 | Abcam, ab185545 (rabbit polyclonal) | 1:1000 dilution | 1:200 (PFA fixed cells) | Developmental cell 56:260-276.e7 |
| Lamp1 | Abcam, ab24170 (rabbit polyclonal) | n/a | 1:200 on isolated phagosomes | Nature communications 14:8354 |
| β-Actin | Cell signaling, 4970, (mouse monoclonal) | 1:1000 dilution | n/a | Zhen Xu, et al. 2025 |
| Commd1 | Novus biology, 03755, (mouse monoclonal) | 1:1000 dilution | 1:200 dilution | |
| Commd9 | Cacalico (UT693) | 1:1000 dilution | n/a | |
| Ccdc22 | Invitrogen, OTI5C1, (mouse monoclonal) | 1:1000 dilution | 1:200 dilution | |
| Ccdc93 | Proteintech, 20861-1-AP (rabbit polyclonal) | 1:1000 dilution | 1:200 dilution | |

| | | | |
|---|---|---|---|
| Vps35l | Invitrogen PA5-28553,(rabbit polyclonal) | 1:1000 dilution | n/a |
